# Neural cognitive signals during spontaneous movements in the macaque

**DOI:** 10.1101/2022.09.05.506681

**Authors:** Sébastien Tremblay, Camille Testard, Ron W. DiTullio, Jeanne Inchauspé, Michael Petrides

**Affiliations:** Montreal Neurological Institute, McGill University, Montréal, QC, Canada; Department of Neuroscience, University of Pennsylvania, Philadelphia, PA, USA; Doctoral Training Center, University of Oxford, Oxford, UK

## Abstract

The single neuron basis of cognitive processing in primates has mostly been studied in laboratory settings where movements are severely restricted. It is unclear, therefore, how natural movements might affect neural signatures of cognition in the brain. Moreover, studies in mice indicate that body movements, when measured, account for most of the neural dynamics in the cortex. To examine this issue, we recorded from single neuron ensembles in the prefrontal cortex in moving monkeys performing a cognitive task and characterized eyes, head, and body movements using video tracking. Despite significant trial-to-trial movement variability, single neuron tuning could be precisely measured and decision signals accurately decoded on a single-trial basis. Creating or abolishing spontaneous movements through head restraint and task manipulations had no measurable impact on neural responses. However, encoding models showed that uninstructed movements explained as much neural variance as task variables, with most of them aligned to task events. These results demonstrate that cognitive signals in the cortex are robust to natural movements, but also that unmeasured movements are potential confounds in cognitive neurophysiology experiments.

## Introduction

In real-world settings, humans think and move simultaneously. A scientist may be reciting an upcoming talk while walking to a conference room, or a soccer player may be deciding who to pass the ball to while running down the field. Primate neurophysiologists have been studying the single neuron computations that enable such cognitive processes for decades (e.g. memory, decision making)^1–4^. However, for practical reasons, it has been difficult to study the single neuron correlates of decision-making while primates move naturally. That is because most single neuron recording technologies and eye movement tracking systems require strict head stabilization and body movement restraint^5–10^. In a well-controlled experiment in monkeys, it is therefore assumed that no body movements should be generated other than those specifically required by the task.

Thus, cognitive neurophysiology experiments in monkeys usually restrict their tracking to movements specifically instructed by the task, such as a lever press or an eye movement (i.e. ‘instructed movements’)^3,11–14^. However, investigators frequently observe that monkeys performing cognitive tasks do move inside primate restraint chairs, sometimes in a way that correlates with the response about to be performed. This observation resembles fidgeting behaviors in humans, such as eyebrow lifting during memory recall, or head tilting while assessing choice options^15,16^. These superfluous, “uninstructed” movements (i.e. not required by the task) often co-occur with cognition in humans as well as in monkeys^17–19^, but their effect on the single neuron signatures of cognitive processing is unclear.

Recent evidence from mice indicates uninstructed movements should not be ignored when trying to explain neural activity. Musall and colleagues trained mice to perform a cognitive task while video recording body movements co-occurring during task execution^20^. Brain-wide calcium imaging revealed those uninstructed movements accounted for most of the neural variance measured, and that many of those movements aligned with task events. This work might suggest that neural correlates of cognitive processing in primates could be the by-product of task-aligned, uninstructed movements that eluded the measurements of neurophysiologists until now. This work calls for precise measurements of body movements beyond the motor effectors involved in task execution in primates.

We combined chronic multielectrode recordings, video tracking, and analytical methods to assess the effects of movement on the neural signatures of cognitive processing in primates. Three monkeys were trained to perform a classical cognitive task under minimal movement restraint. At all times, monkeys were allowed to move their eyes, head, arms, legs and torso while performing a touchscreen-based task. A head-free eye tracker, head tracker, and a high-resolution camera monitored body movements while neural activity was simultaneously recorded from population of neurons in the caudal prefrontal cortex. To maximize the relative presence of task-related neural signals, the task and brain region were strategically selected based on causal studies showing the necessity of this brain region for task performance.

We found that monkeys exhibited many uninstructed movements during performance of the task beyond the motor response required. However, dynamic changes in body posture and field of view did not erase single neuron tuning for the task and cognitive representations could still be accurately decoded from neural ensemble activity on a single-trial basis. Creating or abolishing uninstructed movements through task manipulations and head fixation did not measurably affect neural population responses. However, encoding models revealed that uninstructed movements explained as much neural variance as task variables, although they did not dominate single-trial dynamics as opposed to what was previously reported in mice. Most uninstructed movements were aligned to task events such that upcoming decisions could be accurately decoded from those spontaneous movements alone. Head fixation had little effect on the decodability of body movements. Taken together, these findings show that neural ensembles maintain a robust cognitive representation during free movement and gaze, opening possibilities to study cognitive neurophysiology in more naturalistic settings in monkeys. However, they also indicate that uninstructed movements modulate neural activity in a high-level cognitive area of the primate brain and thus can be important confounds in cognitive neurophysiology experiments in both highly-controlled and naturalistic settings.

## Results

### Cognitive task performance under minimal movement restraint

Three male macaque monkeys were trained to perform a computerized version of the visual conditional association task used by Petrides^21–23^ in lesion experiments demonstrating the causal necessity of prefrontal area 8A, but not neighboring prefrontal areas, for task performance (**Fig. 1a, b**). Head-free monkeys were seated in a modified primate chair that allowed full body movement, only restricting the ability to jump out of the chair. Before recording, monkeys learned to associate 36 randomly selected visual cues from an image bank with two visual targets presented randomly at 8 possible locations on a touchscreen. A set of four different instruction cues was associated with the same two targets that were presented at each recording session (Monkey K: N = 14, Monkey B: N = 6, Monkey L: N = 12 sessions). The instruction cues predicted the correct response, i.e. the target that should be selected on each trial (e.g. If cue A, select target X). All monkeys were implanted with 96-channel Utah arrays in prefrontal area 8A under sterile surgical conditions (**Fig. 1c**). Each array was custom-designed to match subject-specific local morphology and avoid major vasculature. On average, we recorded from 145 neurons simultaneously during each session, including single and multi-units. The number of recorded neurons per session was stable over the course of the experiment (**Supp. Fig. 1**).

**Figure 1.**
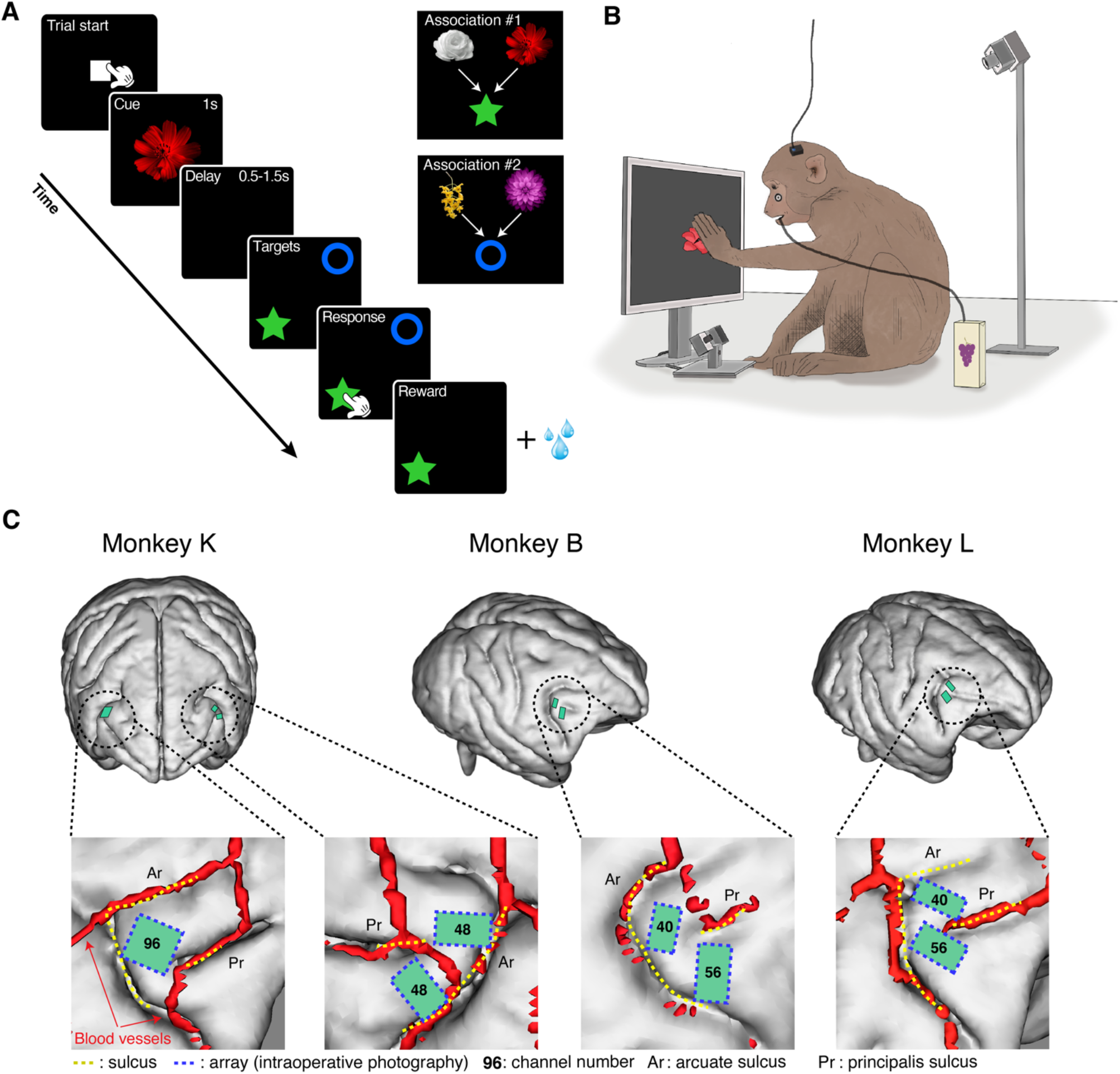
Task design and neural implants. (A) Conditional associative task. Three monkeys previously learnt 36 associations between randomly selected visual instruction cues and two targets. In each session, two pairs of the instruction cues were included, each pair associated with one of two targets. The monkeys initiated a trial by touching a white square on a touch screen with either hand. An instruction cue was presented, followed by a variable delay period. After the delay, two targets were presented randomly at 2 of 8 possible locations. The monkeys were rewarded with fruit juice for selecting the correct target based on the learnt association. (B) Monkeys were seated in a primate chair in front of a 19-inch touchscreen. Monkeys were only restrained at the neck level to prevent them from getting out of the primate chair. Their head, arms, and body were free to move without restraint. Head, arms, hands and tail movements were captured by a video camera, and gaze, head position, and pupil diameter were captured by a head-free tracking system. (C) Implantation surgeries were planned based on 3D reconstructions of brain structure and neuro-vasculature using 3T MRI. Three monkeys were implanted with Utah arrays (green rectangles) within the pre-arcuate gyrus of the prefrontal cortex, where cytoarchitectonic area 8A is found. The exact positions of the implants are depicted based on intra-operative photography of sulcal landmarks (yellow dashed line) aligned to MRI. Numbers indicate channel count on each array. Ar: arcuate sulcus, Pr: sulcus principalis.

Monkeys were trained to expert level on the task before the beginning of the recording sessions (mean hit rate ≥ 90% for all monkeys; **Fig. 2a, Supp. Fig. 2a**). The length of recording sessions was kept short to maintain a constant level of engagement throughout performance of the task (duration ∼1h; **Supp. Fig. 2b**). During task performance, eye position, head position and pupil dilation were measured using a head-free eye tracking system (Eyelink Remote, SR Research). Head-free tracking revealed a significant number of random, uninstructed movements from the eyes and head over the course of a single trial (**Fig. 2b-d, Supp Fig. 3**), and over the course of a typical recording session (**Fig. 2e, f**). Video recording of the monkey using a high-resolution camera (IDS Inc.) installed on top of the primate chair allowed tracking of head, arms, hands, tail, and nose movements during performance of the task (**Fig. 2g**), which also showed a considerable degree of trial-to-trial variability over the course of a recording session (**Fig. 2h, i**).

**Figure 2.**
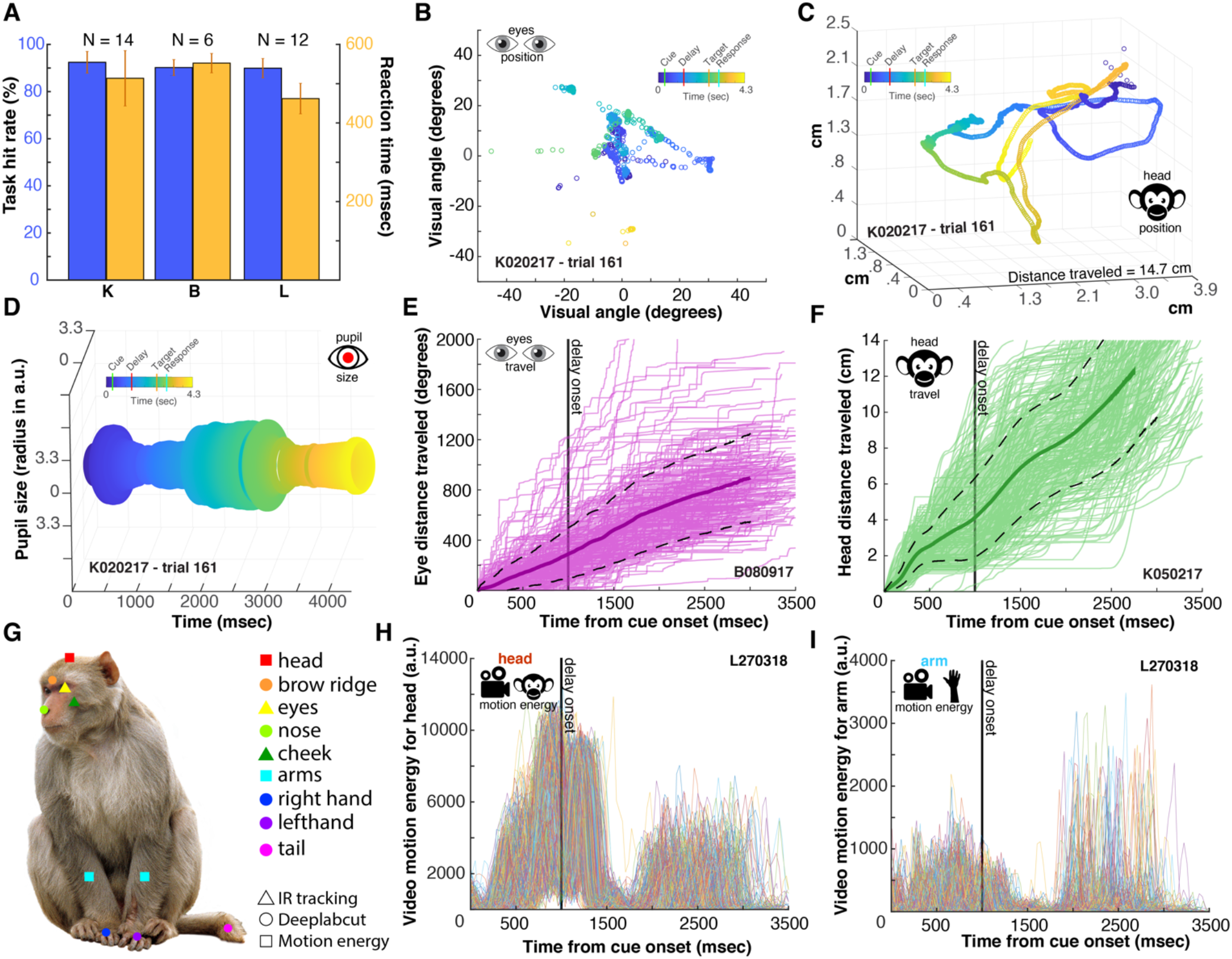
Behavioral performance and movement tracking. (A) Mean hit-rate (blue) and median reaction time (yellow) during the behavioral task for each monkey averaged across sessions. Error bars in A & B are standard deviations (STD). (B) Example trial depicting head-free gaze tracking in a monkey. Inset and colors represent trial time structure. (C) Example trial depicting head position tracking in a monkey. Monkeys were free to move their head during the task. (D) Example trial depicting head-free pupil size tracking over time. (E) Cumulative eye distance traveled (in degrees of visual angle) for each trial (purple lines) of an example session. Monkeys made many uninstructed saccades throughout all trials. (F) Cumulative head distance traveled for each trial (green lines) of an example session. Monkeys made many uninstructed head movements throughout all trials. (G) Video tracking of macaque body movements during task performance. Various cameras and analyses techniques were used to quantify movements of 9 body parts. (H) Head motion energy over time for all trials (colored lines) of an example session. (I) Arm motion energy over time for all trials of an example session.

### Single neuron signatures of cognitive processing during movement

To assess the single neuron signature of cognitive processing during free movement and gaze, we applied standard selectivity analyses to every recorded unit (N=4,414 units, see Methods). Given the cognitive process required by the task (i.e. selecting the particular target associated with particular instruction cues), special attention was given to neuronal selectivity during the delay epoch where no visual stimuli are displayed on the screen and no motor response plan can be prepared (targets appeared at randomly selected locations after the delay). Units that selectively represented the associated target during the delay epoch were named “target-selective”. Selectivity was defined based on a statistical threshold of *P* < .01 comparing mean firing rate across target conditions (one-way ANOVA, over a 500 msec bin, see Methods). An example of a “target-selective” unit can be seen in **Fig. 3a,b**. These units are considered to encode a cognitive representation independent from sensory and motor signals. Units that were selective for instruction cue identity during the cue presentation epoch were labelled “cue-selective”. Finally, units that were selective for the location of the chosen target regardless of the target identity were called “location-selective”. An example of a location-selective unit can be seen in **Fig. 3c** (**Supp. Fig. 4**). Across all recording sessions, we found a large number of each unit type, a result replicated across all three monkeys (**Fig. 3d**). Importantly, these single unit selectivities were observed in the context of large inter-trial variability in gaze and body positions, suggesting the presence of a cognitive decision signal that is robust to movements and changing visual field (see **Fig. 2 & Supp Fig. 3**).

**Figure 3.**
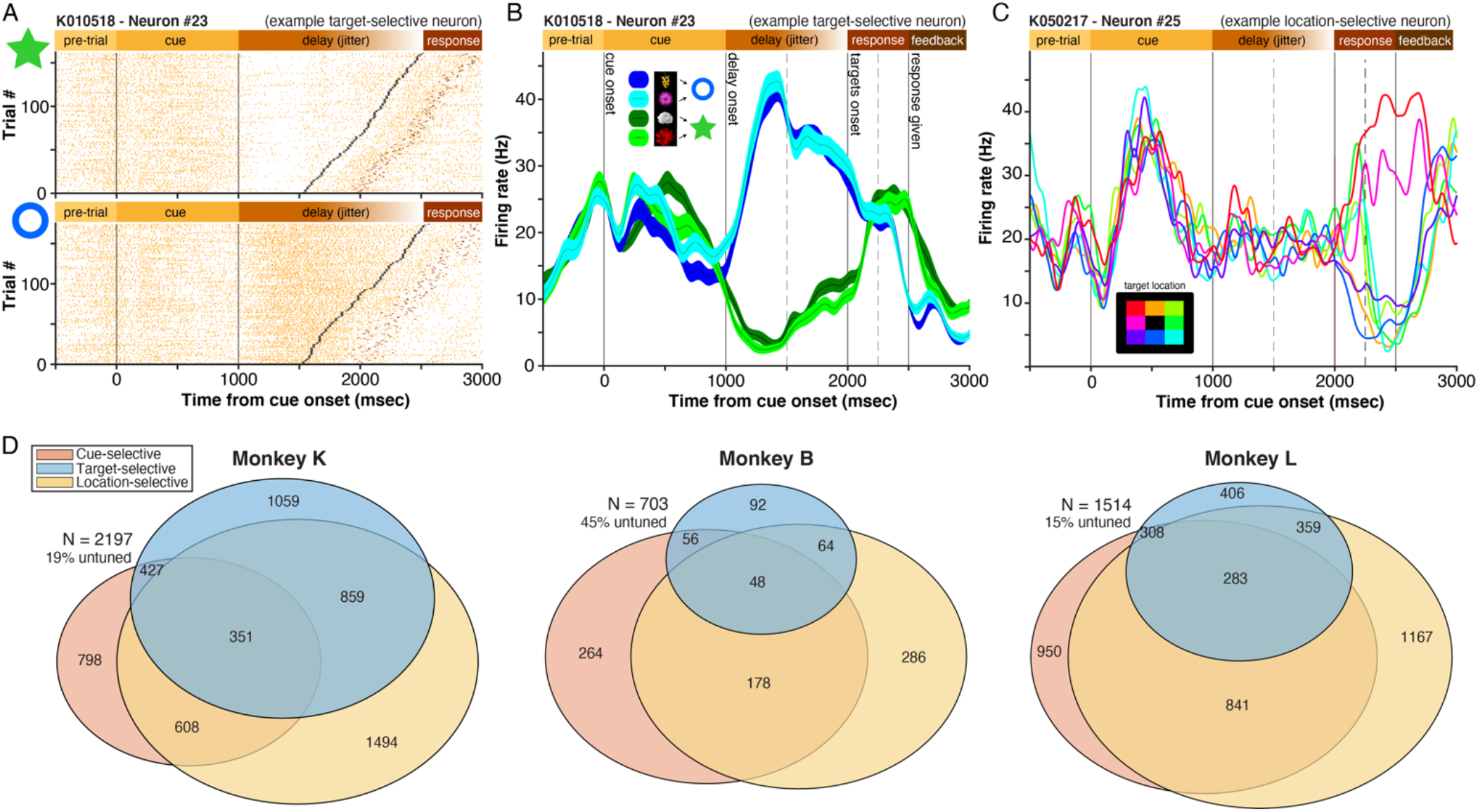
Single neuron and population analyses. (A) Example neuron raster plots for trials where the star (top) or the circle (bottom) was the correct target. Trials are sorted on the basis of the length of the delay epoch, which ends with the onset of targets (black ticks). (B) Example single neuron spike density functions (SDF) showing firing rate as a function of time for each instruction cue presented in one example session (four colored lines). The two blue curves represent trials in which two visually distinct cues were presented, but where the same target (circle) needed to be selected at the end of the trial. Error bars represent SEM. Dashed vertical lines represent cuts in the time axis to account for jitter of the delay length. (C) Example neuron SDF where firing rate is separated based on the spatial location of the correct target (colored curves, 8 locations), regardless of target identity. Selectivity arises only after the onset of the targets. (D) Venn diagrams showing the proportion of each selectivity type across the entire population of recorded units. Selectivity is defined based on specific time epochs (i.e. cue selectivity during the instruction cue epoch, etc., 500msec window size, ANOVA, P < .01). Numbers represent total area or overlap of areas (e.g. in monkey K, 1059 are target selective, 859 are both target selective and location selective).

### Dynamic coding and mixed selectivity during movement

To understand the extent to which units dynamically changed their selectivity over short periods of time during free movement, we conducted a selectivity analysis using a small sliding window of 200 msec (**Fig. 4a**). We looked for the same three types of selectivities described above (instruction cue, target, location) in steps of 100 msec over the length of each trial. A number of cells exhibited cue selectivity about 100-200 msec after cue onset on the screen (red bars) and cue selectivity slowly waned over the course of the delay epoch. Almost simultaneously, a large number of units started encoding the identity of the target associated with the instruction cue presented on that trial (blue bars), many seconds before the target even appeared on the screen. This target selectivity is maintained for the rest of the trial, including the response epoch. Finally, location selective units (yellow bars) were only found during the response epoch, which is expected since the location of the chosen target is unknown to the monkey before then. It is worth emphasizing that this strong spatial selectivity was found in a context of significant trial-to-trial variability in body movements and eyes position (see **Fig. 2e, f** & **Supp. Fig. 3**). This observation suggests the presence of a body-centered spatial reference frame, as opposed to the retinotopic reference frame typically described in head-fixed paradigms in this brain region ^24–26^. These results show that single units dynamically encoded key task variables predicted by the results of the lesion studies conducted in this brain area in unrestrained monkeys ^21–23^. They also show that head fixation may bias interpretation of neural activity towards lower reference frames (e.g. retinotopic).

**Figure 4.**
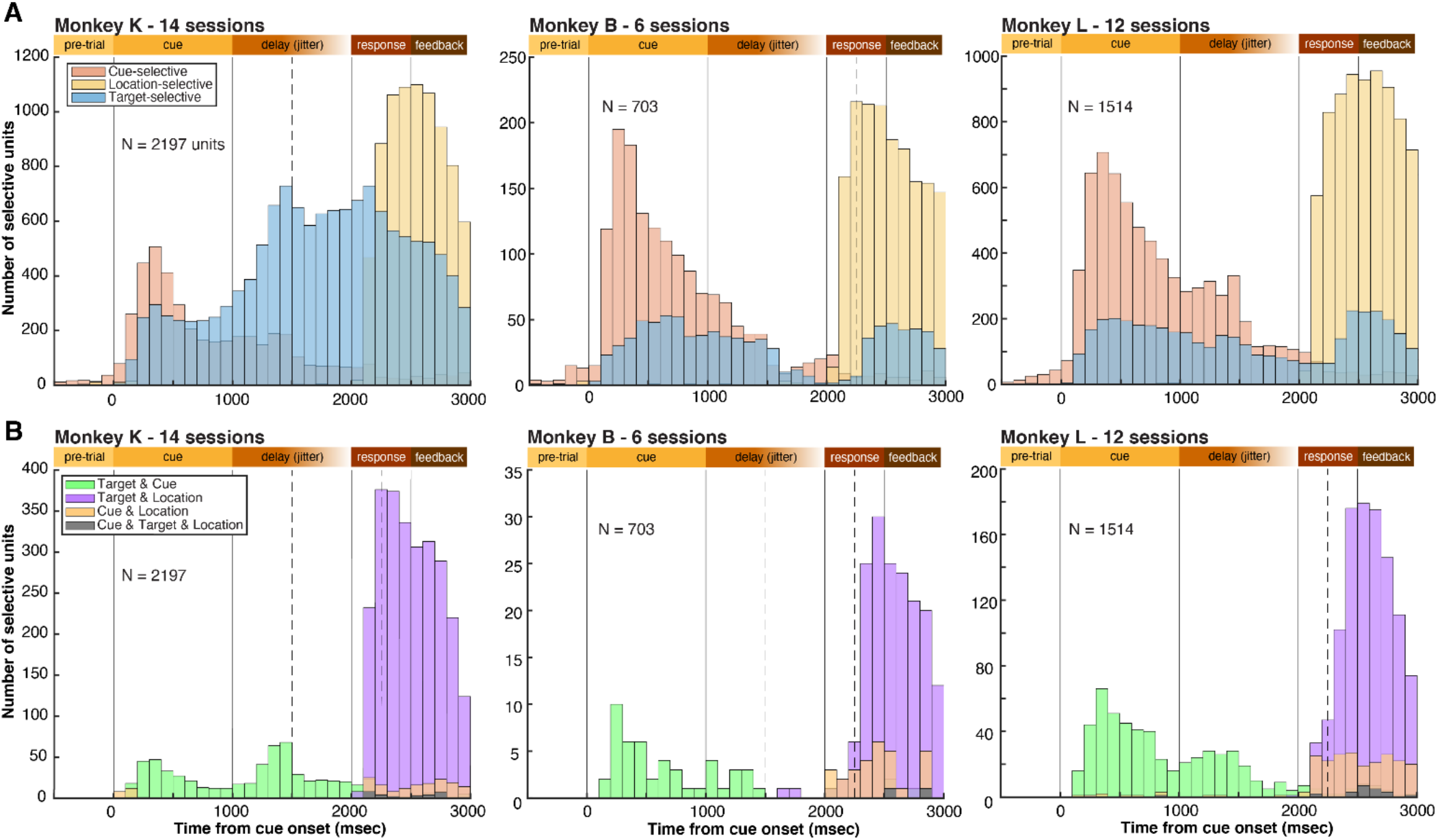
Single neuron and population analyses. (A) Histograms showing the number of instruction cue-, target-, and location-selective units for each time bin of all trials (ANOVA selectivity threshold at *P* < .001, 200 msec bin size). All units from all sessions are included (notice varying Y-axis across monkeys). “N” represents the sum of all units recorded across all sessions for each monkey. (B) Histograms showing the number of mixed selective units for each 200 msec time bin over trials; figure conventions as in (A) (e.g. “Target & Location” indicates that the neuron is both selective for the target and the instruction cue simultaneously in this time bin).

With selectivities dynamically changing over time, we looked for evidence of mixed selectivity, whereby a single unit represents multiple variables simultaneously in its neural firing. We looked at four possible combinations: cue & target selectivity, cue & location selectivity, target & location selectivity, and cue & target & location selectivity. The same 200 msec sliding window with 100 msec steps was used to look for mixed selective signals. In all three monkeys, a large number of units showed mixed target & location selectivity during the response epoch (purple bars; **Fig. 4b**). These were neurons that fired in preference when a given target identity was presented at a given location on the screen. These units representing this specific combination of features may be critical in transforming a cognitive signal (target identity to be chosen) into a body-centered spatial coordinate (target location) that can guide action during natural movement. Mixed selectivity was prevalent despite dynamic changes in body position and visual field over the course of single trials.

### Single-trial decoding and neural ensemble representations

Although averages over multiple trials are informative, the brain cannot benefit from this noise reduction method when generating natural behavior on a moment-by-moment basis. The information content of the neural code on a single-trial basis is therefore a more realistic metric. We used a single-trial decoding approach selecting only neural ensembles of simultaneously recorded neurons to assess the information content of the neural code during free gaze and movement. Replicating a previous approach^25^, we used a support vector machine (SVM) algorithm to decode either the instruction cue identity, the target identity, or the target location from instantaneous neural ensemble activity within a sliding window of 400msec over trials (libSVM, 5-fold cross-validation, see Methods). **Figure 5** plots the instantaneous decoding accuracy for these three variables, averaged across sessions, for each monkey separately (for monkeys combined and interhemispheric comparisons, see **Supp. Fig. 5**). The chance levels are deducted from the number of possible classes (cue: 4, target: 2, location: 8) and confirmed with shuffled label controls (**Supp. Fig. 5b)**. Insets compare decoding accuracies in key task epochs to control data where the labels are shuffled during training. In all three monkeys, instruction cue, target, and location could be decoded with above-chance accuracies on a single-trial basis despite simultaneous free movement and gaze (*t*-tests, *P*<.001, FDR corrected). Importantly, target identity (blue line) could be decoded during the delay epoch when no stimuli were on the screen and no motor response could be prepared, indicating that the conditional associative process that causally depends on this brain region could be decoded from this neural population on a single-trial basis during free, spontaneous movement and gaze. We conclude from this result that cognitive signals in this area are “robust” to natural movements, meaning that they are detectable despite a dynamically changing sensory and motor environment.

**Figure 5.**
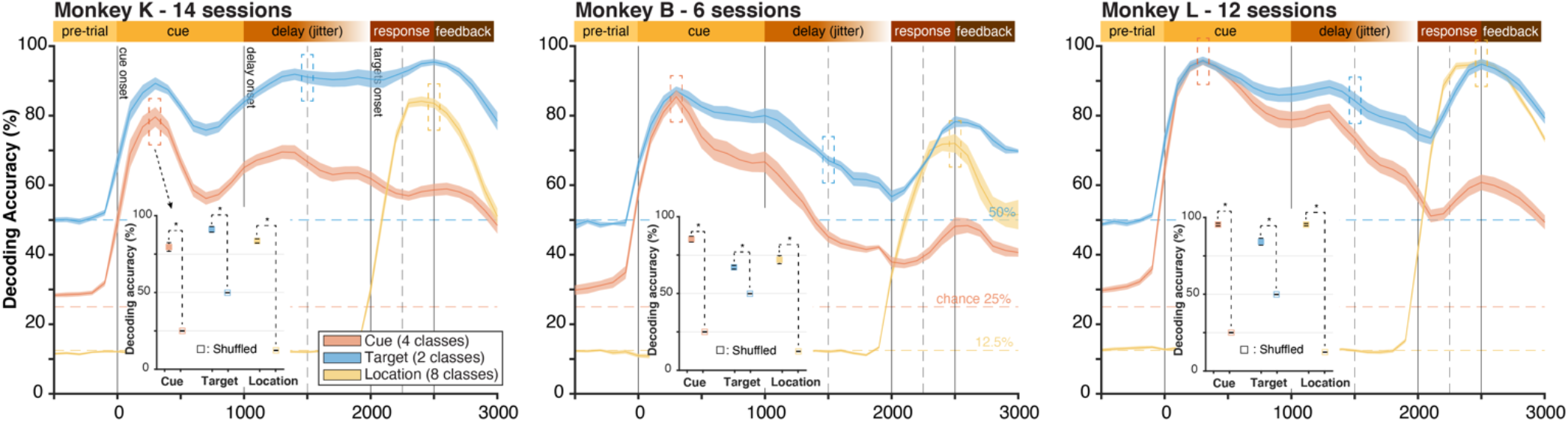
Decoding analyses from neural ensembles. Decoding accuracies of support vector machine (SVM) classifiers predicting cue identity (red), target identity (blue), or target location (yellow), from the firing activity of simultaneously recorded neural ensembles (one ensemble per session). The SVM are trained and tested on a running time window using a bin of 400 msec sliding by 100 msec steps. Horizontal dashed lines represent chance level for each of the three classifications. Insets represent decoding accuracies at key trial epochs (400 msec window) compared to control conditions where training labels are shuffled. *: *P* < .001, FDR corrected. Error bars represent SEM.

### Manipulating uninstructed movements

Although cognitive processing can be decoded from neural ensemble activity during free movement, it is possible that uninstructed movements that are aligned and correlated with task variables could be an important confounding source of information. If, for example, a subject tilts the head one way when about to choose option A, and the other way for option B, the movement itself could drive the neural activity and be mistaken for task-related cognitive activity. In one of the monkeys (monkey K), such a stereotypical movement was observed by the authors whereby a gaze and head position bias during the delay epoch revealed the option to be chosen (**Fig. 6a,b**). To assess the contribution of this movement bias to neural activity, we conducted three control tasks: 1) we modified the task to abolish this bias in monkey K, 2) we modified the task to create a similar bias in a monkey that had no detectable prior bias (monkey L), and 3) we conducted additional sessions in both monkeys with head-fixation to restrain head movements. Modified tasks and head-fixed sessions were not recorded in monkey B.

**Figure 6.**
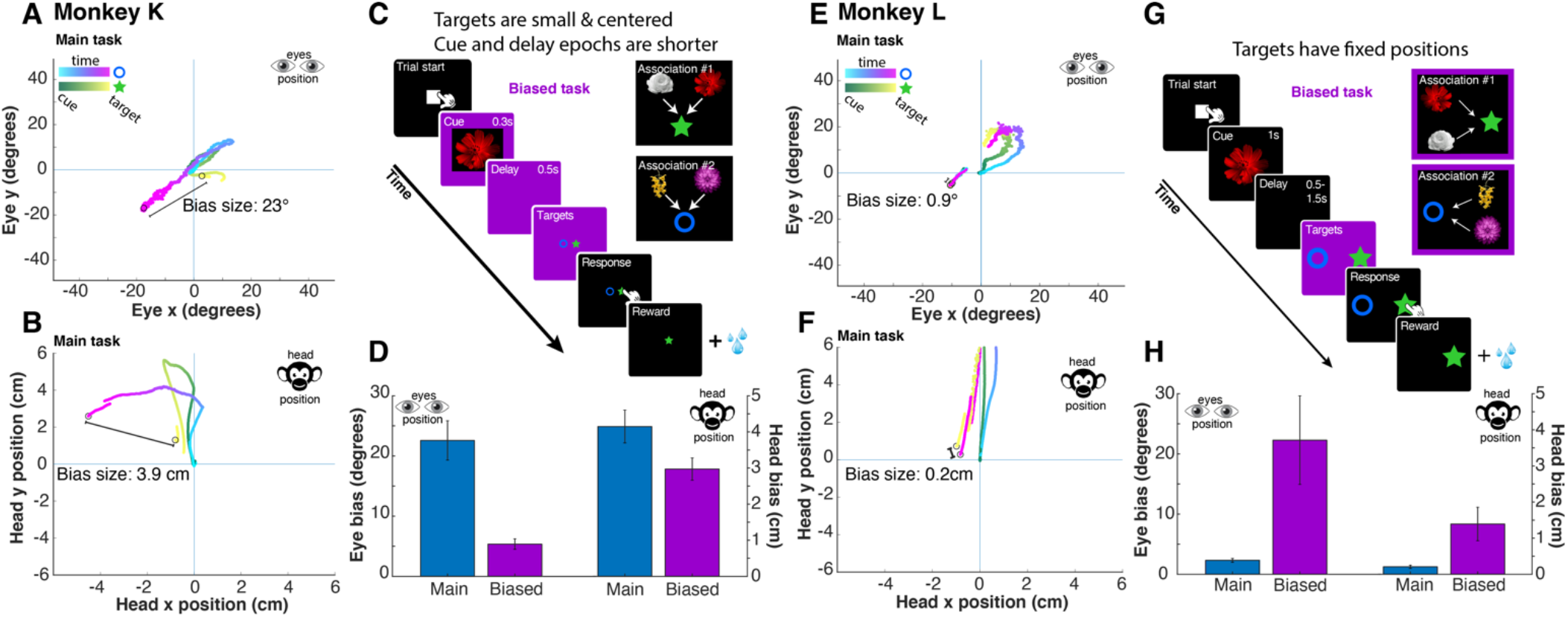
Manipulation of uninstructed movements through modified tasks. (A) Monkey K developed a superstitious movement bias whereby he would position his gaze and head (B) to the left during the delay epoch when he was about to choose the circle, and the opposite if he was about to choose the star. (C) Modified task used to abolish this superstitious bias in Monkey K. (D) Reduction of eye and head bias in modified ‘Biased’ task compared to main task. (E) Monkey L did not have a prior bias in eye or head position (F) during the delay. (G) Modified task with fixed target positions to induce a bias in monkey L similar to the one seen in monkey K. (H) Successful induction of a similar eye position bias during the delay epoch in monkey L following task modifications.

The movement bias in monkey K was controlled by shortening the length of the delay epoch and positioning the targets within a small radius at the center of the screen (**Fig. 6c**). This led the monkey to focus attention quickly on the center screen and did not allow for a stereotypical movement to occur, as shown by a 4.3-fold reduction in spatial bias in eye position (**Fig. 6d**). In monkey L who did not have a movement bias (**Fig. 6e,f**), we modified the task such that targets would appear at fixed locations, i.e. star on the right, and circle on the left (**Fig. 6g**). This led monkey L to develop a spatial bias during the delay epoch similar to the one observed in monkey K (23° bias in monkey K vs 22° in monkey L) (**Fig 6h**). Finally, head fixation using a face shield during additional sessions restricted head movement in both monkey K and L compared to head free sessions (−83% and −62% in mean head motion energy for monkey K and L, respectively; **Supp. Fig 6**).

We ran a new set of analyses with these modified tasks to examine if the reduction or creation of these uninstructed, task-aligned movements would have an influence on neural population responses. As with the rest of the study, only results replicated across monkeys are discussed. **Figure 7** compares the neural population tuning for the main task (‘Main’), the head restrained task (‘Fixed’), and the modified tasks (‘Biased’). Note that for monkey K’s biased task, a stereotypical movement was reduced, whereas for monkey L, such a movement was created. When comparing head free to head fixed sessions in both monkeys, we did not observe a significant increase or reduction in the proportion of neurons tuned to the instruction cue, target, or location variables, despite a large reduction in head movement (**Fig. 7a**, Chi-square test, *P* > .05, FDR corrected). The same was true for mixed selectivity in both monkeys (**Fig. 7b**), and for single-trial decoding accuracy of these variables from neural ensembles (**Fig. 7c** & **Supp. Fig. 7**). In addition, in monkey K, reducing stereotypical, task-aligned movements (‘Biased’) did not influence the accuracy of single-trial decoding or population tuning, even for target identity, with which the movement was highly correlated. Similarly, in monkey L, the creation of a movement bias that correlates with a cognitive task variable (‘target identity’) did not lead to a noticeable increase in decoding accuracy or population tuning for this variable. We also didn’t observe differences in the strength of neural tuning across the three behavioral manipulations in either monkey when using eta-squared metrics or receiver operating characteristics curves (**Supp. Fig. 8**). In summary, these findings indicate that manipulating uninstructed head and eye movements that are highly correlated with task variables does not impact the single neuron tuning measured in this brain area.

**Figure 7.**
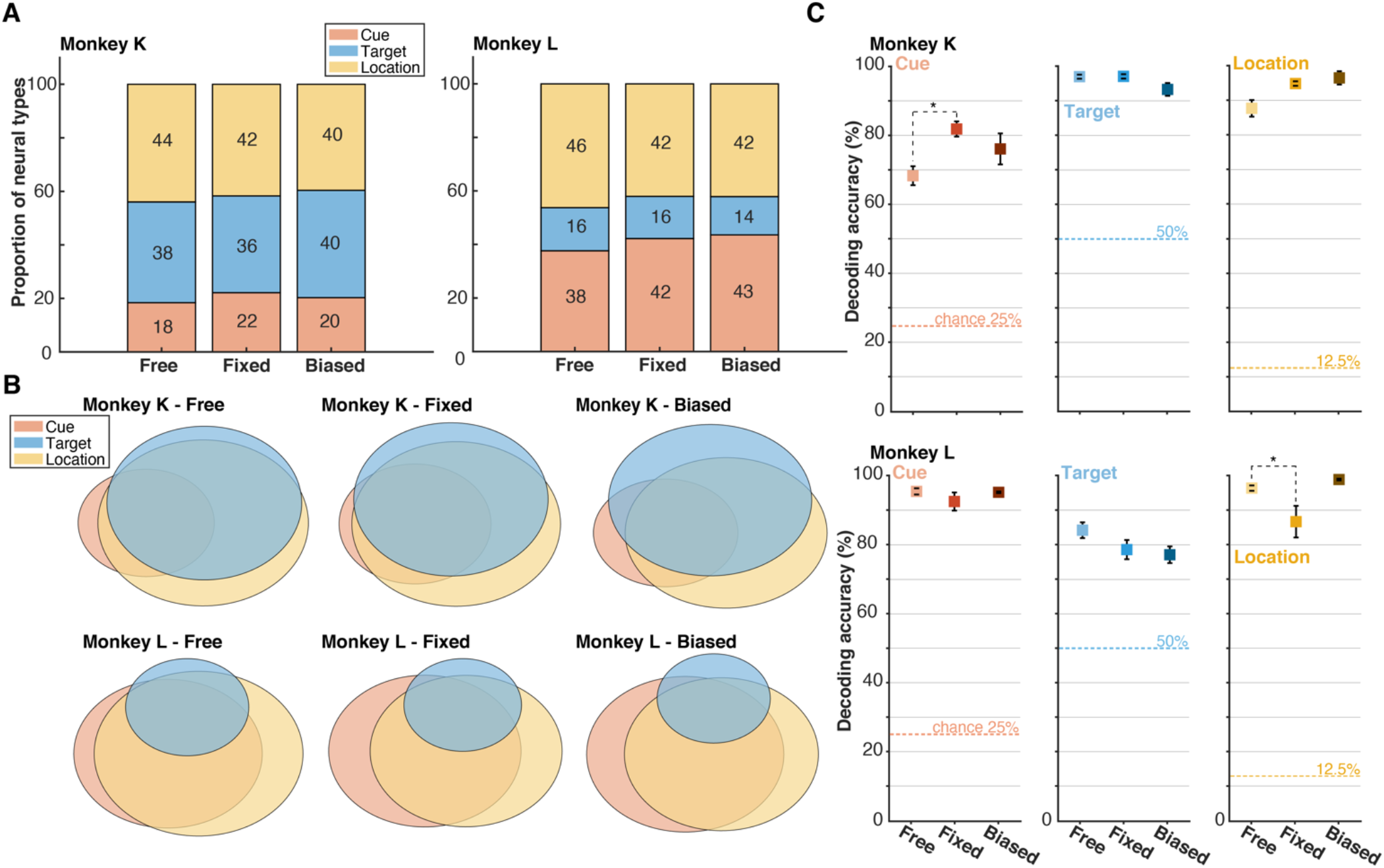
Modified tasks have no effect on neural population tuning. (A) Effect of head fixation (Fixed), induced movement bias or reduced movement bias (Biased) compared to main task (Free) on the proportion of neurons tuned to task variables instruction cue, target, and location. (B) Same as in A, but also showing the mixed selective neurons across tasks, as represented by the overlap between circles, as in **Fig. 3d**. (C) Single-trial decoding accuracies for the three task variables, as in **Fig. 5**, but comparing across modified tasks. *: *P* < .05, FDR corrected. Error bars represent SEM.

### Decoding cognition from uninstructed movements

While an obvious head movement bias was detected by the authors in monkey K (**Fig. 6A, B**), we wondered if other motor effectors (arms, hands, nose, brow ridge, tail, etc.) exhibited any bias that correlated with the decision variable. We used a similar decoding approach as above when decoding task variables from neural activity (libSVM, see Methods), however, here we asked whether the decision variable could be decoded from uninstructed body movements alone during the delay epoch, that is, before any motor response can be planned. **Supp. Fig. 9a** reveals that many motor effectors carried information about the upcoming decision, such that the target to be selected could be predicted from the position of the head, face and limbs during the delay epoch (Monkey K: 95%, Monkey L 70% accuracy from all body parts combined). This was possible even in Monkey L, whom had no apparent movement bias. We note important differences between the two monkeys, which suggests that inter-individual differences exist in how much body movements assist decision making in primates. Importantly, we find that head-fixation, despite significantly reducing the ability to move the head (−83% and −62% in head motion energy for monkey K and L, respectively), had little impact on the predictive power of residual body movements (Monkey K: 95% vs 96% for head-free vs. head-fixed, respectively; Monkey L: 70% vs 68%; **Supp. Fig. 9b**). This suggests that movements that are typically not restrained or tracked in monkey experiments can correlate with decision variables on a single-trial basis, potentially driving observed cognitive signals.

### Neural encoding of task and movement variables

We used a multivariate encoding approach to delineate how much of the neural population responses were respectively explained by task variables and potentially confounding movement variables. Adapting code provided by Musall and colleagues^20^, we closely replicated their methodology to quantify the total and unique neural variance explained by each movement and task variable, as well as groups of variables classified as task, instructed movements, and uninstructed movements (see **Supp. Table 1** for all predictors). In brief, we built a linear encoding model (i.e. ridge regression) that used a large array of task and movement variables to predict the neural activity of ensembles of simultaneously recorded neurons on a single-trial basis (see Methods for details). The model design matrix included discrete event variables (e.g. monkey’s choice) as well as continuous, analog variables (e.g. arm position) (**Fig. 8a**, left). Analog variables were converted to additional discrete events by using thresholding. The continuous signals were fit using a single linear scaling coefficient. For discrete event variables (e.g. a screen touch), we produced a time-varying event kernel relating the specific event to neural ensemble activity (**Fig. 8a**, middle). The time-varying kernel accounted for potentially shifted neural responses to the event, associating a model coefficient to every time-shifted copy of the event. A modeled neural response was generated for each recorded neuron at each time point (every 33 msec) over the course of a recording session based on all predictors and model coefficients (**Fig. 8a**, right).

**Figure 8.**
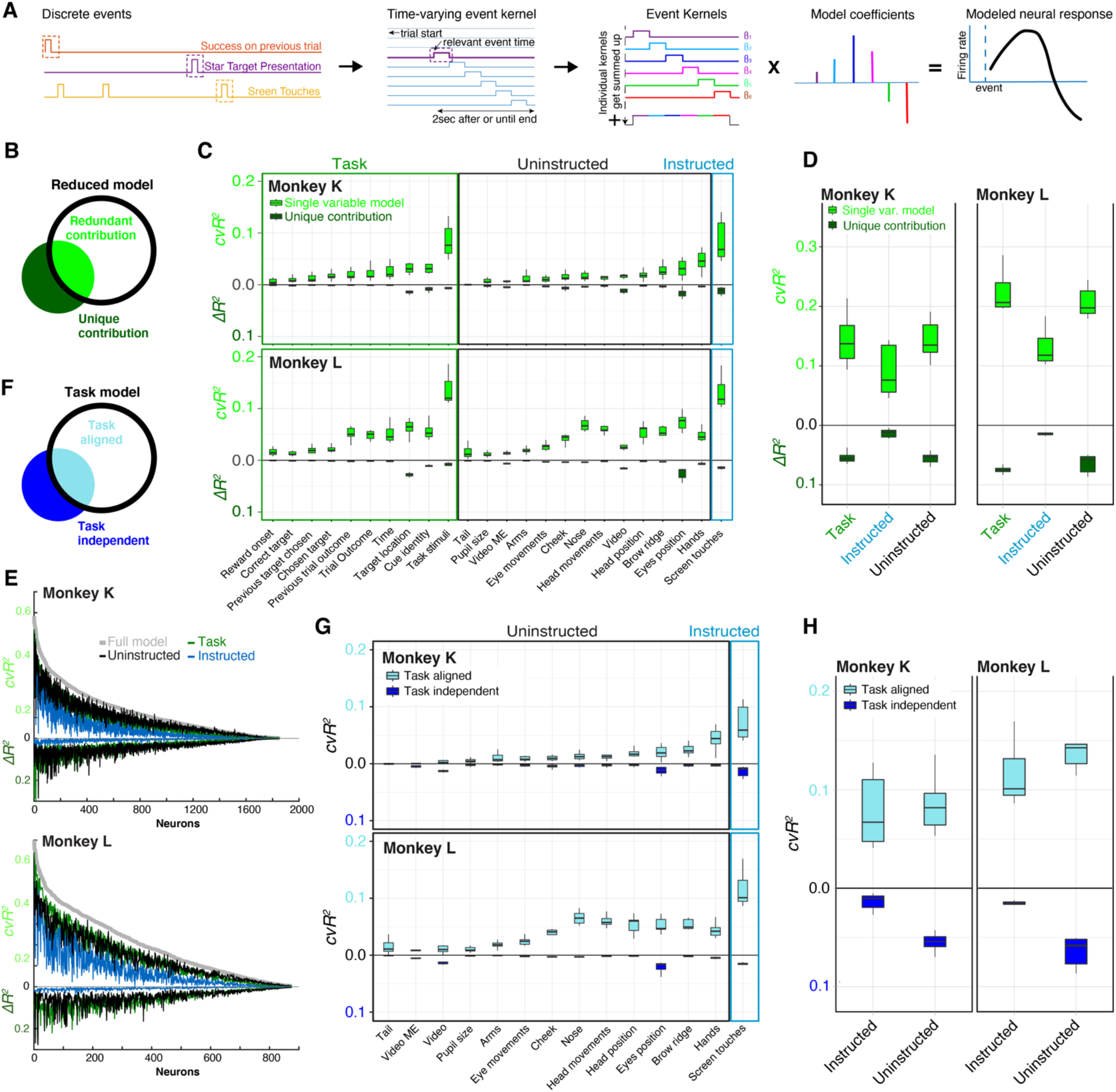
Neural encoding of task, instructed, and uninstructed movements. (A) Discrete task and movement events as well as analog, continuous variables (B) are included in a linear encoding model (ridge regression). A 2SD threshold is applied to extract discrete events from analog movement signals. (C) A single scaling coefficient is applied to continuous variables, and (D) time-shifted copies of discrete events are created to account for delayed neural responses. (E) A coefficient is estimated for each time varying event kernels to model neural responses to events on a single trial basis. (F) The contribution of each variable to the model is divided into its total contribution (light green), which may have a redundancy with other model variables, and its unique contribution (dark green) that excludes potential redundancy. (G) Full (light green) and unique (dark green) contribution of each model variables to neural activity predictions in both monkeys. (H) Full and unique contributions from variable groups for both monkeys. (I) Full and unique contribution from variable groups for each single neuron recorded in the entire experiment.

To assess model performance in predicting neural ensemble activity, we computed the tenfold cross-validated *R^2^* (*cvR^2^*). The cross-validation ensures that the variance explained is not the result of overfitting despite a high number of model variables. We also shuffled neural activity to test the validity of the model predictions (shuffled control: *cvR^2^* < 2e10^−5^ in all recordings). The full model predicted on average 21.27% and 30.65% of all neural variance in monkey K and monkey L, respectively, which is higher than results previously obtained in mice using multielectrode probes (≈10% *cvR^2^*)^20^. Next, we looked at the individual contribution of each model variable to understand which one contributes the most to the neural activity predictions. First, we computed models shuffling all variables except one variable of interest and did so iteratively for every model variable (‘Single variable model’) (**Fig. 8b**, light green). This provides an upper bound on the variance explained by each model variable. Because information provided by a given variable can be redundant with other model variables, we also compared the accuracy of the full model to the one of a reduced model where we shuffled only the variable of interest (‘Unique contribution’) (**Fig. 8b**, dark green). This provided a lower bound on the amount of information contributed by this variable.

**Figure 8c** depicts the model accuracy (*cvR^2^*) for each single variable model as well as the unique contributions in terms of difference from the full model (Δ*R^2^*). In both monkeys, task stimuli, which include cue and target onset/offset, explained the most variance (Full contribution: 8.5% for monkey K and 13.6% for monkey L), on par with touchscreen responses (Full contribution: 8.34% in K, 13.05% in L). Eye position had the largest unique contribution (mean = 1.57% for monkey K and 2.6% for monkey L). Overall, we did not find that movement variables made significantly larger contributions than task variables, as opposed to the dominance of movement variables observed by Musall et al (2019) in mice. Variables such as ‘Video’ and video motion energy (‘Video ME’), which explained a large portion of the variance in the cited study, explained a similar amount of variance to task variables such as cue identity or trial outcome in the present study in monkeys (see Methods for discussion on the relation between encoding and previous decoding analyses).

To compare the overall contribution of instructed or uninstructed movements relative to task variables, we reran the analysis using groups that included all variables pertaining to task (21 predictors), instructed movements (3 predictors), or uninstructed movements (31 predictors) (**Fig. 8d** & **Supp. Table 1**). Surprisingly, we found that task and uninstructed movements had similar explained variance in both monkeys (Monkey K: task 14.32%, uninstructed 14.11%; Monkey L: task 22.38%, uninstructed 20.7%). Instructed movements had slightly lower explained variance (Monkey K: 9.06%, Monkey L: 13.04%). When looking at unique contributions, task and uninstructed movements also had similar explained variance (Monkey K: uninstructed 5.65%, task 5.46%; Monkey L: uninstructed 6.5%, task 7.49%). When looking at single neurons individually, we found that the majority of neurons replicated this pattern, with neurons equally representing task variables and uninstructed movements variables in their spiking activity, followed by instructed movements (**Fig. 8e**). This held true both for full contribution (top traces), and unique contribution to model predictions (lower traces).

Although movements did not dominate single trial dynamics, they did explain as much neural variance as tasks variables that are the focus of cognitive experiments in monkeys. We wondered whether these movements could be potential confounds by measuring the proportion of neural activity explained by movements that are “task aligned” vs “task independent” (see Methods). Here, “task-independent” refers to the explained variance gained by adding a given movement predictor on top of a model including all task variables. On the other hand, “task-aligned” contributions refer to the difference between the full explanatory power of a given movement variable (**Fig. 8c**, light green) and its task-independent contribution. Looking at each motor effector individually, we find that most have little task-independent contributions and relatively larger task-aligned contributions to neural activity (**Fig. 8g**). When grouping effectors in “instructed” and “uninstructed” movement categories, we find that instructed movements have a small task-independent and a large task-aligned contributions, which is expected considering those specific movements were required by the task (**Fig. 8h**). However, we find that uninstructed movements have a large task-aligned contribution, even larger than task-independent (Monkey K: independent 5.64%, aligned 8.47%; Monkey L: independent 6.47%, aligned 14.23%), meaning that most uninstructed movements were performed in relation to some task variable (**Fig. 8h**). We also find that the movement variance of most effectors can be explained by task events alone, even when those task events are restricted to the main variables of the task (cue, target, and location identity) (**Supp. Fig. 10**). This indicates that a large part of the neural variance is explained by spontaneous movements uninstructed by the task in monkeys, and that those movements are highly correlated to task variables for which neural tuning is typically computed in cognitive experiments.

## Discussion

First, the results demonstrate that a cognitive decision signal is robustly represented in the primate cortex during naturalistic movements and gaze. Standard single-neuron activity metrics, such as tuning and mixed selectivity, could be reliably extracted and neural ensemble representations could be decoded on a single-trial basis despite large variability in sensory-motor environment. However, we found that many spontaneous body movements were performed in correlation with the task and the upcoming decisions. When modeling 31 body movement variables on a single-trial basis, including both those instructed and uninstructed by the task, we found that movements explained as much neural variance as task variables. Although we did not find that movement variables dominated cortical activity, as observed in mice^20,27^, we found that most of these movements were aligned to specific task events and cognitive variables, potentially confounding cognitive neurophysiology experiments in nonhuman primates.

Traditionally, the neural correlates of cognition in the nonhuman primate brain have been studied in highly controlled experimental settings where movements are physically restricted by means of primate chairs and cranial implants ^6,7,10^. In such paradigms, a single trial is aborted if an uninstructed movement occurs, such as an eye movement away from the fixation point^13,25,28,29^. Consequently, it remains unknown how spontaneous eye, head and body movements modulate neural activity in higher brain areas underlying cognitive processing. In the current study, we demonstrate that decision signals were robust to natural movements in a prefrontal area that had been demonstrated to be causally related to the cognitive processing examined ^21–23^. Despite this causal dependency, a large part of the neural activity was explained by uninstructed movements that were task-aligned, suggesting that movements not required by the task should be measured during cognitive task performance in order to disentangle cognitive signals from spontaneous movement signals in monkeys. This observation is of critical importance for the growing field of neuroethology that strives to study more naturalistic, unconstrained behaviors in primates ^30–32^. Since head-fixation did not prevent uninstructed movements from aligning with task events, we believe our findings also apply to more controlled laboratory experiments where these movements are typically unmonitored.

Our study differs from that of Musall et al.^20^ in important aspects which may explain some discrepancies in the amount of neural variance explained by movements. First, we intentionally targeted a single brain area (prefrontal area 8A) that is required for the cognitive task selected, as opposed to the cortex-wide survey afforded by the wide-field calcium imaging technique used in mice. By doing so, we believe to have maximized the potential presence of task-related signals in the neural population studied. Finding the correct match between task and brain area may be crucial in estimating the relative prevalence of cognitive signals, and mismatches may lead to overestimating the relative contribution of movement variables. A second important difference with the study by Musall and colleagues is the animal model under study. Mice and primates diverged **~**100 million years ago, such that findings from mice research sometimes do not translate to nonhuman primates^33,34^. One interpretation of the Musall et al. findings is that cognition relies on neural machinery that originally evolved for movement, repurposing algorithms fine-tuned for movement to engage in cognition^35,36^. Perhaps in nonhuman primates, association cortical areas evolved neural machinery independent from movement circuits, such that cognitive signals can be fully dissociated from the immediate sensory-motor environment^37–39^.

Finally, prefrontal area 8A has been largely described by neurophysiologists to be involved in the planning and control of eye movements^25,40–42^. Although a relatively large degree of neural variance was explained by eye movements in the present study, other effectors, such as the hands and head, were also represented. From these correlational results alone, one might conclude that area 8A is involved in head movement, or hand reaches, although lesion studies demonstrate no deficit in the planning and control of head, hand, and even eye movements after complete bilateral ablation of this brain area^21–23,43^. This raises the question: what proportion of neural signals we are measuring that really are efference copies, or other forms of modulation, with no causal role in the behavior examined? The observation of movement signals in the prefrontal cortex does not, in our view, indicate a causal involvement of this cortex in performing uninstructed movements. Thus, what is the role of these movement signals, and why are they as prevalent as signals related to the cognitive function this brain area implements? Why did our manipulations of uninstructed movements not alter the neural representation of task variables? We argue that there is a need for primate neurophysiologists to carry out causal investigations of neural activity during natural behavior to understand the contribution of movement signals to cognitive processing in real-world settings.

## Methods

### Subjects

Three adult male cynomolgus monkeys (macaca fascicularis) participated in the experiments: monkey K: 7 kg, monkey B: 8 kg, monkey L: 7 kg. All procedures complied with the requirements of the Canadian Council of Animal Care and the Montreal Neurological Institute animal care committee. Over the course of a testing session (once a day), the monkeys would receive their daily amount of fluids consisting of water-diluted fruit juice for correctly performing the task. In addition, the monkeys would receive daily fresh fruits and vegetables at the end of each recording session. Each session lasted on average one hour, and no more than 1.5 hour. Physical and mental health was assessed daily by veterinary and laboratory staff throughout the course of the experiment. No animals were sacrificed for the purpose of this study.

### Surgical procedures

Surgical plans were prepared with the help of brain MR images obtained on a Siemens 3T scanner (TIM TRIO, Montreal Neurological Institute). T1-weighted images (MP-RAGE) with and without gadolinium enhancement (Gadovist®, Bayer, Germany) were obtained to reconstruct the 3D cortical surface and cerebral vasculature (Brainsight Vet 2.4, Rogue Research, Canada; see **Fig. 1c**). Custom Utah arrays (Blackrock Microsystems, UT) were designed on the basis of the local morphology in the region of interest for each monkey. The aim was to cover optimally the region of interest while avoiding major blood vessels. All surgical operations were carried out under isoflurane general anesthesia and under strict sterile conditions with the help of experienced veterinary staff continuously monitoring vital signs. The animal head was positioned in a stereotaxic frame (Kopf Instruments, CA) and a midline skin incision was made to expose the dorsal aspect of the skull. The temporalis muscles were retracted ventrally and a square-shaped craniotomy (1.5 × 1.5cm) was made in the fronto-lateral bone, based on MRI coordinates, with a dental drill equipped with a diamond round-cutting burr (Horico, Germany). The dura-mater was exposed and a dural flap was performed extending ventrally. Direct visualization of cortical landmarks, such as the arcuate sulcus and the sulcus principalis, permitted identification of the pre-arcuate convexity where area 8A of the prefrontal cortex lies in the macaque brain^43,44^. The array(s) were carefully positioned over the region of interest and implanted using a pneumatic inserter held by a flexible surgical arm. The dural flap was closed with 5-0 Vycril sutures and a layer of dura regeneration matrix (Durepair, Medtronic, MN) was laid over the reconstructed flap. The bone flap was thinned with a drill and replaced over the craniotomy and secured to the skull with low-profile titanium plates and screws (DePuy Synthes, IN). The Cereport connector was secured caudally to the skull opening using eight 1.5mm diameter titanium screws with a length determined by the skull thickness as measured by pre-operative MRI. The two reference wires were inserted between the dura and the cortex and the exposed portion of those wires and of the array wire bundle were coated with a thin layer of Quick-Sil (WPI, FL) or Geristore (Denmat, CA). The muscle, fascia and skin were closed in anatomical layers with absorbable 3-0 Vycril sutures and the Cereport connector protruded through a small opening in the skin. The monkeys recovered for two weeks before the first recording session. No headposts were implanted for this study and no acrylic or dental cement was used in surgeries.

### Behavioral task

In the current study, the monkeys were trained to sit in primate chairs that did not restrict their head or arm movements. The front panel of the chair was removed allowing the monkeys to reach outside the chair with their arms. The primate chair did not impose head movement restrictions, allowing the monkeys to turn around in the chair and look in any direction (360°). The monkeys were positioned in front of a 19-inch touchscreen (ELO touch 1937l, Accutouch, CA) connected to a behavioral control computer running MonkeyLogic 2.0 (version 47, NIMH, Hwang et al. 2019) on a Windows 7 PC. The chair was positioned 7 inches from the touchscreen, such that the monkeys could reach easily at all locations on the screen. At this distance, the screen occupied 86.4 degrees of visual angle horizontally and 73.7 degrees of visual angle vertically.

The three monkeys were trained on a computerized adaptation of the conditional association task used in Petrides ^21–23^ (**Fig. 1a**). In this previous study, monkeys with bilateral lesions of prefrontal area 8A (the area targeted in the current study) exhibited specific impairments on this task, while sparing cognitive performance on other equally difficult tasks, such as working memory tasks. A trial was initiated by the monkey touching a white square appearing at the center of the touchscreen. Following the touch, one out of four possible instruction cues was shown on the screen for 1 second. After cue presentation, a delay period of random duration (0.5-1.5 sec) followed and then two target stimuli appeared at random locations. Note that there were 8 possible locations and the stimuli always appeared opposite to each other. The monkeys received a reward when they touched the correct target stimulus, i.e. the target that was predicted by the instruction cue presented a few seconds earlier in the trial. Upon selection of the correct target, the untouched target disappeared to give feedback on what target was selected, and a squirt of juice was delivered through a metal straw (Crist Instrument, MD). Before the beginning of this experiment and over the course of multiple training sessions, the monkeys learned 36 arbitrary cue-target associations, always using the same two target stimuli (a blue circle and a green star), until a threshold of 80% correct choices was reached. In any given recording session (one session per day), 4 cues were selected (2 associated with the circle, 2 associated with the star) and randomly interleaved from trial to trial. In some recording sessions, the cues were interleaved in a block design, whereby a first pair of cues (1 associated with each target) was presented for half the session, and a second pair for the other half of the session. The latter block design was used for 32 recording sessions (Monkey K: 14, Monkey B: 6, Monkey L: 12) and is referred to as the “main task”. The design whereby the 4 cues are randomly interleaved from trial to trial was used for 8 recording sessions (Monkey K: 5, Monkey L: 3). It was thought that the latter task would be more difficult for the monkeys because 4 cues had to be maintained “online” at all time to perform the task. In fact, there was no difference in performance between blocked and interleaved designs (data not shown).

Behavioral performance at the task was calculated using a hit rate (correct trials / correct trials + error trials) and the median reaction time was computed for correct trials (**Fig. 2a**). Hit rate and reaction time was also computed separately for each of the 8 possible target locations to identify potential spatial biases (**Supp. Fig. 2a**). Hit rate over time was also computed to look for signs of fatigue or satiation in the monkeys during a recording session (**Supp. Fig. 2b**). All trials in a recording session were split in three thirds based on their chronological order, and a separate hit rate was computed for each third of the trials.

For a better understanding of the effects of body movements on neural activity, we designed a third version of the task, referred to as the “biased task”, in which spatial biases were induced by modifying target locations in a non-random manner (i.e. associating a target with a specific position). This created spatial expectations for target locations and induced stereotypical movements during the delay epoch. Alternatively, we also created a biased task whereby existing, spontaneously generated biases were extinguished by similar procedures involving the manipulation of target locations. This allowed the parsing out of the individual contribution of those uninstructed movements to neural activity.

Although the monkeys were free to move their head and arms in most sessions, we also ran a series of sessions with the monkeys’ head fixed to the primate chair (referred to as the “head-fixed” sessions). Since the monkeys were not implanted with head posts, we fixed the monkey’s head using a solid, transparent face shield resting on the monkey’s snout and attached to the chair (see **Supp. Fig. 5a,b**). The face shield did not prevent the monkey to move his eyes or arms. It only limited the ability of the monkey to move his head and turn around in the chair. This manipulation was useful to understand the contribution of head movements specifically to neural activity and to make a more direct comparison to more conventional neurophysiology experiments conducted with head-fixed monkeys. The performance and reaction time for the “head-fixed” and “biased task” was calculated using the same method detailed for the “main task”.

### Behavioral and video monitoring

All behavioral monitoring devices were synchronized to the master clock of the neural recording system. All touchscreen touches by the monkeys were recorded by MonkeyLogic and marked as discrete events. In addition, we used a head-free eye tracking system to monitor eye position, pupil size, and head position (in 3D) with a 500 Hz temporal resolution (Eyelink 1000, Remote version, SR Research, Canada) (**Fig. 2c-d**). This was achieved with the help of a small sticker put on the monkeys’ right cheek at the beginning of each session (see **Supp. Fig. 5a**). This eye tracking signal was lost when the monkeys made very large head movements (e.g. turning 180° in the chair) bringing the sticker out of the sight of the camera. Additionally, we video-recorded the body movements of the monkeys using a video camera installed atop the primate chair (IDS UI-3240ML-C-HQ, 30 fps, Germany). The camera provided a good view of the monkeys’ head and arms but did not capture potential leg and torso movements occurring inside the primate chair (**Supp. Fig. 5a** and **Fig. 2g**).

Using the head-free eye-tracker, we computed the cumulative distance traveled by the gaze on the screen over time for each trial in units of visual angle (degrees). This was obtained by calculating the Euclidean distance between each eye sample point in a trial (**Fig. 2e**). The same computation was done on the head movement samples to calculate the cumulative head distance traveled over trials, this time using the Euclidean distance in 3D space (cm) (**Fig. 2f**). Using the eye and head-tracking data, we looked for potential biases in eye or head movements that could correlate with the decision the animal made in each trial (i.e. to select the star or the circle target). To do that, we calculated the trial-averaged eye or head position during the delay epoch, and separated trials per target selected (star or circle). The distance between the average positions for star-selected and circle-selected trials is referred to as “spatial bias” and indicate that different uninstructed movements are made as a function of the decision being taken (see **Fig. 6a,b**).

Using the data from the video camera attached to the primate chair, we computed the motion energy for the arms and head during performance of the task. The arms and head were parsed using manually drawn regions of interests (ROI) in the Motion Energy Analysis software (MEA v4.10, F. Ramseyer 2019, https://osf.io/gkzs3/). Motion energy was computed as the number of pixels that “change” value within the ROI on a frame by frame basis. A “change” was defined based on a threshold crossing operation on the pixel color change to eliminate video noise (threshold = 8 a.u.). Motion energy over time for each trial was plotted for the arms and head for an example session (**Fig. 2h,i**). The motion energy for the head was compared across sessions of the “main-task” with the monkeys’ head free to move and sessions of the “head-fixed” task (**Supp. Fig. 5c,d**). A large stereotypical head movement of Monkey L at delay onset was captured by the motion energy, but missed by the Eyelink camera because the movement was so large that the camera lost tracking of the reference sticker.

### Pose estimation using DeepLabCut

For body part tracking we used DeepLabCut (version 2.2) [Mathis et al, 2018, Nath et al, 2019]. Specifically, we labeled 200 frames taken from 3 videos/animals (then 95% was used for training). We used a ResNet-50-based neural network with default parameters for 150,000 training iterations. We validated with 10 number of shuffles, and found the test error was: 6 pixels, train: 307,200 pixels (image size was 640 by 480). We then used a p-cutoff of 0.6 to condition the X,Y coordinates for future analysis. This network was then used to analyze videos from similar experimental settings. An example labeled video per animal can be found in the online Methods.

### Neural recordings and spike detection

The neural data from the chronically implanted Utah arrays was recorded using a Cereplex Direct 96-channel neural recording system (Blackrock Microsystems, UT). The raw signal was bandpass filtered (0.3Hz to 7.5KHz) and digitized (16 bits) at 30,000 samples per second by a Cereplex E digital headstage installed on the Cereport connector of the Utah array. The digitized signal was routed from the headstage to the recording computer (PC, Windows 7) using a long, flexible micro-HDMI cable that did not impede the movements of the monkeys. We also collected neural data using the Cereplex W wireless system from Blackrock Microsystems but were overall better satisfied with the quality of the data obtained with the wired Cereplex E solution. For each channel, neural action potentials (or “spikes”) were detected online based on a channel-specific voltage threshold equal to 4 times the root mean square of the noise amplitude (Central Suite software, Blackrock Microsystems). The waveforms and timestamps of each spike were saved to disk and transferred to Offline Sorter (version 2.8.8, Plexon Inc., TX) for manual sorting of spike waveforms. Well-isolated single units as well as multi-unit clusters were classified on each channel and saved for later analyses in Matlab (MathWorks, MA). On average, we recorded from 171 (SD 12.7) units (including single and multiunits) from Monkey K on each session, 130 (SD 7.2) from monkey B, and 135 (SD 11.3) from monkey L. We made no assumptions as to whether the recorded units were the same or different ones from session to session. The number of recorded neurons as a function of sessions (in chronological order) is plotted in Supp. Fig. 2A and was stable for all three monkeys over the course of the experiment.

### Single neuron selectivity

Each neuron was characterized through a series of analyses revealing its selectivity to different task parameters. For each of the 4,414 units recorded throughout the experiment, we outputted 15 figures characterizing the “profile” of the unit consisting of: 1) its anatomical location on the array(s), 2) its waveform, 3) its raster plot (1 kHz resolution) separated by target type (trials in which the star was the target vs. trials in which the circle was the target). Trials were ordered by delay length. 4) same as #3, but with trials ordered by chronological order of presentation. This provided a view of low-frequency time-varying responses over the session, such as those correlating with block changes in the main task. 5) Spike-density functions obtained by smoothing each trial’s raster using a 45-msec gaussian kernel and averaging across trials with the same target (star vs. circle). 6) Same as #5, but grouping trials based on instruction cue identity rather than target (4 cues per session). 7) Same as #5, but grouping trials based on the spatial location of the target (8 locations). 8) “Running effect size” analyses, whereby the instantaneous effect size for instruction cue, target, and location was calculated at each 100 msec of the trial and plotted as a smooth curve over time. The effect size was calculated using the eta-squared derived from an ANOVA comparing the mean response across trials grouped by instruction cue, target, or location, using a sliding window of 200 msec. This allowed us to see how selectivity changed rapidly over short timescales, and how mixed selectivity arose during trials. 9) A bar plot showing the response at target onset (+/− 150 msec), grouped by location (8 locations/bars). An ANOVA was computed on those 8 means to determine spatial selectivity at target onset. 10) Same as #9 but focusing on the response epoch (when the monkey touches the screen) using a +/− 75 msec window centered on the response. 11) “Neural drift” analyses examined slow changes in firing rate over the course of a session (hereby named “drift”). This drift was calculated by sliding a 4-minute window by steps of 1 minute over the course of the entire session, regardless of trial type. The average firing rate was computed within each window and plotted as a smooth curve over time (typically one hour). This permitted detection of the presence of slow, large fluctuations in firing rate that are the result of changing isolation of the unit during recording (e.g. when an electrode slowly drifts away from a neuron and its spikes become progressively merged with noise). Because of the block design of our main task, such drift can artificially inflate the selectivity of units to parameters that change across blocks, such as cues (see below). 12) Same as #5 but corrected for drift, 13) same as #6 but corrected for drift, 14) same as #7 but corrected for drift, 15) same as #8 but corrected for drift.

Upon inspection, neural drift was observed on some of the recorded units. Drift is problematic in a block design task because any task parameter that changes on a timescale similar to the drift will appear to be “coded” by the neural activity. For example, if a neuron’s isolation is lost around the middle of the session, its firing rate will be null for the second half of the session, leading to a large “selectivity” if any spikes were present during the first half of the session. On the other hand, drift will also artificially increase neural response variability over the session, underestimating the true coding accuracy of any task parameters that do not change with blocks (such as target or location selectivity).

We therefore implemented the following drift correction policy to calculate accurate neural coding values:

1. Eliminate neurons that have a large drift, as defined by a 25% change in average firing rate between the first half and the second half of the session.
2. Exclude sessions that have more than 15% of their neurons exhibiting a large drift, as defined in step 1.
3. Drift correct all remaining neurons by subtracting baseline firing rate from instantaneous firing rate. As a result of step 2, 1/15 sessions was eliminated for monkey K, 4/10 sessions were eliminated for monkey B, and 1/13 sessions was eliminated for monkey L. In step 3, the objective was to eliminate any residual effects of neural drift by baseline-correcting the instantaneous firing rates of each unit. The baseline firing rate was calculated using a 4-minute sliding window, as defined above for analysis #11. The firing rate recorded during each trial was subtracted by the baseline firing rate, leading to a mean of 0 Hz.

### Population selectivity

In population analyses, the set of all neural units recorded across sessions was pooled into a single group, or population for each monkey. That corresponds to 2197 units in monkey K, 703 units in monkey B, and 1514 units in monkey L. To characterize the proportion of each neural type in the population, we classified units based on their selectivity for the instruction cue, the target, or the location of the target. “Cue-selective units” were identified based on the firing rate of the unit 100 to 600 msec after instruction cue onset, separated by cue condition (4 conditions per session). An ANOVA compared the mean firing rate across trials of the same cue condition and a statistical threshold of *P* < .01 was used to define a selective unit. “Target-selective units” were identified using the same procedure by focusing on the epoch during the delay 100 to 600 msec after cue offset. “Location-selective units” were identified using the epoch 250 msec before to 250 msec after the touch response was delivered on the touchscreen. All three epochs were 500 msec long. Venn diagrams were generated to display the relative proportion of each neural type using the *venn* function for Matlab (©Darik Gamble, 2009).

In order to look at the selectivity of the units evolving over time, we computed additional analyses of each unit’s selectivity at all the time points of the trial. For this analysis, we computed selectivity for instruction cue, target and location using a 200 msec sliding window with steps of 100 msec. At each step, an ANOVA was computed comparing the firing rate across conditions (cue: 4 conditions, target: 2 conditions, location: 8 conditions), as detailed above. For each time point, a unit was considered selective at that time point if it passed a threshold of *P* < .001. The number of units being selective for the cue, the target, or the location of the target at each time point was cumulated and plotted on a peri-stimulus time histogram showing the total number of units in the population exhibiting a given selectivity at a given time. Importantly, a unit could be selective for more than one task feature at the same time point.

Many neural units exhibited mixed selectivity, that is, selectivity for multiple task features at the same time. We expanded the previous time-dependent analyses to include exclusively units that were selective for a combination of features in a given 200 msec time window. We classified units into “cue & target”, “target & location”, “cue & location”, or “cue & target & location” selective units for each time point. The same *P* < .001 statistical threshold was used. The number of mixed selective units at each time point over the trial was plotted as a peri-stimulus time histogram.

### Decoding analyses

We ran a series of decoding analyses using a machine-learning algorithm to estimate the amount of information contained in the neural firing code over time. In those analyses, we used a support vector machine (libSVM^45^) with 5-fold cross-validation. For each SVM, we only included simultaneously recorded units, hereby named “neural ensembles”, to get a more realistic estimate of information content on a msec time resolution and to better account for confounding factors that can affect coding, such as noise correlations^25,46^. For each cross-validation fold, non-overlapping training and testing sets of trials were defined, and the accuracy of the trained model was calculated based on the number of correct predictions on the testing set (correct predictions / all predictions). The number of observations per class (cue: 4 classes, target: 2 classes, location: 8 classes) was balanced using random sampling so that each class had the same number of observations as the smallest class. To account for the random sampling process, we ran 20 iterations of each SVM and computed the mean decoding accuracy across those iterations. Chance performance of the decoder was obtained by randomly permuting the labels (shuffled control) before training and following the same analysis procedure.

We ran multiple decoders to extract information pertaining to the instruction cue, the target, or the location of the target based on the instantaneous firing rate of neural ensembles. For each of these three decoding analyses, we ran the decoder on a sliding window to obtain the decoding accuracy at each time point of the trial and observe dynamic changes in information over time. Each decoder was run on a 400 msec time window with all simultaneously recorded neurons in a recording session as features, and all trials in that session as observations. The window was then stepped by 100 msec and the decoding analysis was repeated using the firing rates in this new window. The sliding process was repeated until the end of the trial. This sliding window decoding analysis was carried out for each session individually, and the average over all sessions for each monkey was computed and plotted in **Fig. 5**. Standard errors of the mean are represented by the shaded area around the average line. Specific time points of interest (cue epoch: 300 msec after cue onset, delay epoch: middle of delay, and response epoch: response onset) were selected for statistical testing of decoding accuracy. Two-sample t-tests comparing the mean decoding accuracy to the mean shuffled decoding accuracy were carried out at each of these 3 time points, and the family of comparisons for the three monkeys (9 tests) were corrected using false discovery rate (Benjamini & Hochberg, corrected *P* < .05).

We ran the same decoding analyses on variations of the main behavioral task (**Fig. 7**). Namely, we were interested in seeing whether fixing the head of the monkeys, rather than letting it move freely, would have an impact on neural activity (aka. head-fixed sessions). We also investigated whether creating or abolishing stereotypical movements through task manipulations would have an impact on the neural code (aka. biased sessions). Only monkeys K and L participated in those variations of the task. In monkey K, those additional sessions were acquired while recording from the right hemisphere array only. Therefore, we compared those additional sessions to main task sessions also acquired from the right hemisphere array only. This explains why results vary slightly for monkey K between **Fig. 6** and **Fig. 7**. Main task (**Fig. 6** also contains main task sessions from left hemisphere array). Decoding accuracies were calculated using the same technique detailed above, using a 400 msec sliding-window SVM with 5-fold cross-validation. Differences in mean accuracy across variations of the task were statistically tested at the same three time points of interest during the instruction cue, delay, and response epoch. Since the “biased” version of the task implied shorter trial lengths in monkey K, average curves were aligned to those time points of interest for plotting purposes (**Fig. 7**, dashed boxes and insets). Two sample t-tests comparing the decoding accuracy during the main task to the ones of each variation of the task were computed using the same P-value threshold as above corrected for multiple comparisons using false discovery rate (Benjamini & Hochberg, corrected *P* < .05). We considered a result to be statistically significant if it crossed the corrected *P* threshold and was replicated across monkeys.

We ran additional decoding analyses predicting the upcoming decision (star or circle target) based off uninstructed movements (**Supp Fig. 9**). We did so by replicating the analysis method above, with the difference that movements we used as predictors rather than neural activity. We used either each motor effectors individually (head, cheek, eyes, left hand, right hand, nose, brow ridge, tail, total head motion energy, total arm motion energy), or all of them together (full body) as predictors in the SVM. Chance level is at 50%. We focused on a window at the end of the delay epoch (500 msec), before the targets onset so the correct movement pattern cannot be planned yet. Therefore, this decoding is based solely on uninstructed, spontaneous movements made by the monkeys.

### Encoding analyses

#### Linear encoding model

We used the linear encoding model methods described in Musall et al. 2019 to determine how much of the variance in neural activity could be explained by task-related variables versus movement variables. We used the code provided by the authors and adapted their model for use with our data to perform the same variance decomposition. We briefly summarize the rationale and details of the model here and refer the reader to the original publication for more details.

The model uses ridge regression to determine the linear relationship between the full array of regressors (i.e. design matrix) and neural activity. Once ridge regression has found the best fitting linear model between the design matrix and the neural activity, we can compare the variance explained by different groups of regressors by manipulating the design matrix and evaluating the effects on model fit. We note here the distinction between “variables” and “regressors”: variables refer to the named variables enumerated in Suppl. Table 1, while regressors refer to columns of the design matrix. There can be multiple regressors associated with any single variable. For example, a variable that is associated with discrete events (e.g. cue presented) can have multiple regressors to capture potential delays between the event and its effect on the neural activity (see section Design Matrix below for more details). In our analyses, variables and their associated regressors were put into one of three variable groups: task variables (e.g. animal’s choice, reward, stimulus presentation), instructed movement variables (i.e. movements by the motor effector required by the task) and uninstructed movements variables (i.e. movements from any other motor effector captured in this experiment). In our task, subjects interacted with a touch screen to perform a cognitive task, thus, screen touches were considered instructed movements. Uninstructed movements accordingly described all other movements which were detected either through the Remote EyeLink tracker (for head, eye, and pupil tracking) or through video camera. Our model contained a total of 44 explanatory variables (See **Suppl. Table 1** for a full table describing each model variable). We use these variables to predict trial-averaged and single-trial neural activity of each neuron recorded simultaneously during a session.

Regularization is needed to provide a more stable estimate of regressor coefficient due to the large number of regressors used in the model. This need for regularization is why ridge regression is used instead of a standard linear regression. In brief, ridge regression enforces an L2 regularization on the regressor coefficients (i.e. the beta values). This regularization drives regressor coefficients for regressors that are not explaining much variance towards zero, and thus helps avoid issues of overfitting. We emphasize the fact that the L2 regularization does not set regressor coefficients to exactly zero to contrast it with another common type of regularized linear regression, Lasso regression. Lasso regression enforces a sparseness prior by using L1 regularization on the regressor coefficients which ultimately results in the model performing feature selection (i.e. eliminating certain regressors from the model by setting their coefficients to exactly zero). This elimination of regressors would introduce challenges when comparing model fits and thus was not used.

The model with regularization can be summarized in the following equations:

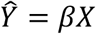

Where X is the design matrix containing all regressors, Beta is the collection of all regressor coefficients, Y_hat is the neural activity predicted by the model with those specific betas. The loss function for the model is:

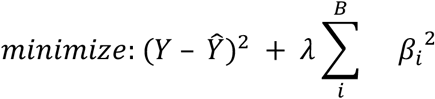

Where Y is the neural activity for each trial during a recording session, and the final term of the equation is the regularization term enforced by ridge regression. For X, Y, and Y_hat, we use the column vector convention such that each row is a time point and each column is a particular regressor.

#### Design Matrix

The model’s design matrix included regressors for two broad types of variables: discrete behavioral events and continuous (analog) variables. Discrete behavioral events included task-related features, such as the animal’s choice and screen touches (**Fig. 8a**). All uninstructed movement variables (except video data, see below) were both a continuous signal (analog) and discrete events in the model. Discrete events for uninstructed movements were captured through threshold detection (threshold: change in variable X is greater than 2SD) (**Fig. 8b**). All variables were downsampled by a factor of 30 such that each bin corresponds to 30.33 ms to match with the frame rate of the video data collected.

Neural activity may have a delayed or multiphasic temporal relationship to discrete events (either task or movement related) and thus our model had to be able to capture such relationships. Following the methods of Musall *et al*. 2019, we utilized time-varying event kernels (**Fig. 8d**). Briefly, each event variable was represented as a set of binary vectors (i.e. each element of the vector takes on a value of zero or one). The set consisted of a vector with a value of one at the time of the relevant event and numerous time-shifted copies of this original vector (**Fig. 8d**). Importantly, each vector of this set has its own regressor coefficient allowing for the model to capture flexibly the neural response at many time points relative to the event (**Fig. 8e**).

Three types of time-varying kernels were defined according to the expected dynamics of neural responses: 1) whole trial – time-shifted copies spanned all frames of a trial, used for events that could impact neural activity during the entire trial (e.g. success on the previous trial); 2) post-event – from event time until the end of the trial; and 3) peri-event – from 1sec before the event to 2sec afterwards (or until the end of the trial if less than 2sec remained). Type 2 kernels were used for task events for which we did not expect neural encoding prior to the event (e.g. target location). Type 3 kernels were used for movement events where we expected preparatory neural activity.

Finally, the model also included analog variables for uninstructed movements (e.g. video). We used Singular Value Decomposition (SVD) to compute the 200 highest-variance dimensions of video data from the animal’s upper body. SVD was performed either on the raw video data (‘Video’) or the absolute temporal derivative (motion energy; ‘Video ME’). Importantly, video frames were cut to exclude the touch screen, such that task events could not be inferred from the video. For each analog variable, ridge regression produced a single scaling weight, making these variables useful in capturing neural responses that scale linearly with a given variable (**Fig. 8c**). As in Musall et al. 2019, we did not use any lagged versions of analog variables, and thus did not account for delayed or multiphasic responses of analog signals.

#### Model fit evaluation: Full contribution vs. Unique contribution

We next sought to address which of the task variables, i.e. instructed and uninstructed movement, best predicted the observed neural activity. Predictions of neural activity were obtained using tenfold cross-validation. For each fold, 90% of the data of a recording session were used to fit the model (i.e. estimate the regressor coefficient for all regressors). This model was then used to predict the neural activity for the held out 10% of the data. The predictions from all folds were combined to produce the cross-validated estimate of neural activity for the entire recording session. The correlation between the cross-validated estimates of neural activity and the recorded neural activity was used to calculate the variance explained by the model. To compute all explained variance by a group of model variables (cv*R*^2^ or full contribution), we created reduced models where all variables apart from the specified ones were shuffled in time. The explained variance by each reduced model shows the maximum potential predictive power of the corresponding group of model variables. To measure each variable’s *unique* contribution to the model, we created reduced models in which *only* the specified variables were shuffled. The difference in explained variance between the full and the reduced model yielded the unique contribution (Δ*R*^2^) of that group of model variables. The unique contribution shows the amount of explained variance in the full model that can be uniquely attributed to that particular group of model variables (i.e. nonredundant to all other model variables, **Fig. 8f**). We compared the importance (cv*R*^2^ and Δ*R*^2^) of all instructed or uninstructed movements relative to task variables using a one-way ANOVA and Tukey post-hoc tests.

#### Task-aligned vs. task-independent contribution

Next, we investigated the extent to which the contributions of movement variables were independent to the task being performed. To compute the ‘task-aligned’ or ‘task-independent’ explained variance for each movement variable group (instructed and uninstructed), we created a ‘task-only’ model where all movement variables were shuffled in time. This task-only model was then compared with other models where only one of the movement variable groups was shuffled. The difference between the task-only model and this second model, with the added contribution of either instructed or uninstructed movement variables, yielded the task-independent contribution of that movement variable group. The task-aligned contribution was computed as the difference between the total variance explained by a given variable group (its cv*R*^2^) and its task-independent contribution.

#### Missing Data Procedure

Because monkeys were not head-restrained, it was possible for them to move in such a way that eye and head tracking from the Eyelink were lost temporarily (e.g., turning around in the chair). We had an average 58% (SD = 10%) of missing data points over all our sessions. To mitigate these data gaps, we performed linear interpolation over short runs (less than or equal to 150 msec) of missing data^47^. With this procedure we were able to recover on average 12% of absent data. The linear encoding model we used does not have a method for dealing with missing data points, accordingly these time points with missing data were removed before fitting the model (mean = 46%, SD = 6%).

#### Linear Encoding Model: Mitigation of multicollinearity

Collinearity between model variables, or multicollinearity when dealing with several variables, reduces the reliability of regressor coefficient estimation in linear regression^48^. Accordingly, it was necessary to take a series of preprocessing steps to reduce multicollinearity in order to have confidence that model performance was accurate. Importantly while these steps do improve confidence that the model performance was accurately measured (i.e. that we have measured the true R^2^ value of the model), they cannot increase model performance (i.e. increase the R^2^ value); only reduce it. We describe these steps below.

The first step of mitigation was to logically eliminate any variables that could be determined from some combination of other variables. For a simple example, consider the variable ‘stimulus location’. There were eight possible locations a stimulus could have in the task, but only seven location variables were included in the model. This was done because the eighth location can be determined by knowing that the stimulus is not present in the other seven locations. Next, we sought to orthogonalize (i.e. mathematically remove multicollinearity) between variables that were likely to be strongly correlated. For example, video variables were very likely to be correlated with other movement model variables, such as head movements detected with the EyeLink tracker, since these movements could likely also be inferred from video data. To accomplish this goal, we used the QR decomposition method described in Musall *et al.* to orthogonalize video variables with respect to all other movement variables and uninstructed movement variables with all instructed movement variables

In our case, we applied QR decomposition to a reduced design matrix, Xr, containing all movement variables which could also be inferred from the video data. As the order of regressors influences how much a particular regressor could be rotated by the QR decomposition, Xr was ordered so that the motion energy and video regressors were the last columns. Since columns 1 to j of *Q* span the same subspace as the columns of the original design matrix, columns of *Q* can be substituted for their respective columns in the design matrix. Accordingly, we replaced the motion energy and video columns of the full design matrix with their orthogonalized columns from Q. This substitution effectively decorrelates the video variables with respect to all other movement variables, allowing the model to use the unique contributions of the video variables to improve model fit while not altering the coefficients of any other variables. Subsequently, we used the same approach to orthogonalize all uninstructed movement variables against instructed movements. This was done to ensure that all information that could be explained by instructed movements could not be accounted for by correlated, uninstructed movement variables. As described above, this only improves our confidence in the accuracy of the fit of the model and cannot increase the explanatory power of uninstructed movement variables.

As a final insurance against multicollinearity between model variables, we utilized a method Musall et al. devised called ‘cumulative subspace angles’ (CSA) on the full design matrix. Briefly, CSA takes advantage of the fact that the R matrix generated by QR decomposition provides a measure of how far each column of a matrix lies from the subspace spanned by the previous columns. Accordingly, this measure can be used to determine the collinearity of each successive regressor (column of design matrix) with all previous regressors. Specifically, CSA generates a collinearity value ranging from zero to one, where zero indicates complete multicollinearity and one indicates no multicollinearity at all Regressors with excessive collinearity (i.e. collinearity values at or very near zero) can be removed. We re-emphasize here the distinction between variables and regressors: removing subsets of regressors is not the same as removing variables. No variable had enough of collinear regressors to be removed entirely from our model. Over all experimental sessions, the median collinearity value was 0.935, indicating a mean angle of 63.45° between each variable and all other variables. For comparison of magnitude, in the original manuscript Musall et al. report a median collinearity value of .84 indicating a mean angle of 57°. From these results, we concluded that we had, as much as possible, mitigated multicollinearity in our design matrix and thus could have confidence in the accuracy of our evaluation of model performance.

### Relationship between tuning and decoding analyses

The encoding analysis mentioned here differs from the decoding analyses and the single neuron selectivities reported earlier in important ways. These differences can help explain why some task variables, such as cue ID, target ID, and location, appear to have lower explanatory power in the encoding model than what would be hinted from the tuning/decoding results. Significant results from tuning and decoding are mostly driven by the trial-averaged difference in firing between different target conditions. Increasing the trial-by-trial variance (with random movements, for example) will reduce the significance of these results only if the variance is so great that the firing rate distributions for each variable level start to overlap. In contrast, significant results from the encoding method are driven by how consistently a variable predicts the exact firing rate response of a neuron on a millisecond basis. This is true for the entire length of the trial with this method, whereas tuning/decoding focuses specifically on the delay epoch. In the case of a categorical variable such as target identity, any deviation from a consistent firing pattern will reduce the explanatory power of the target identity variable in predicting the instantaneous firing rate.

## Supplementary material

**Supp. Figure 1.**
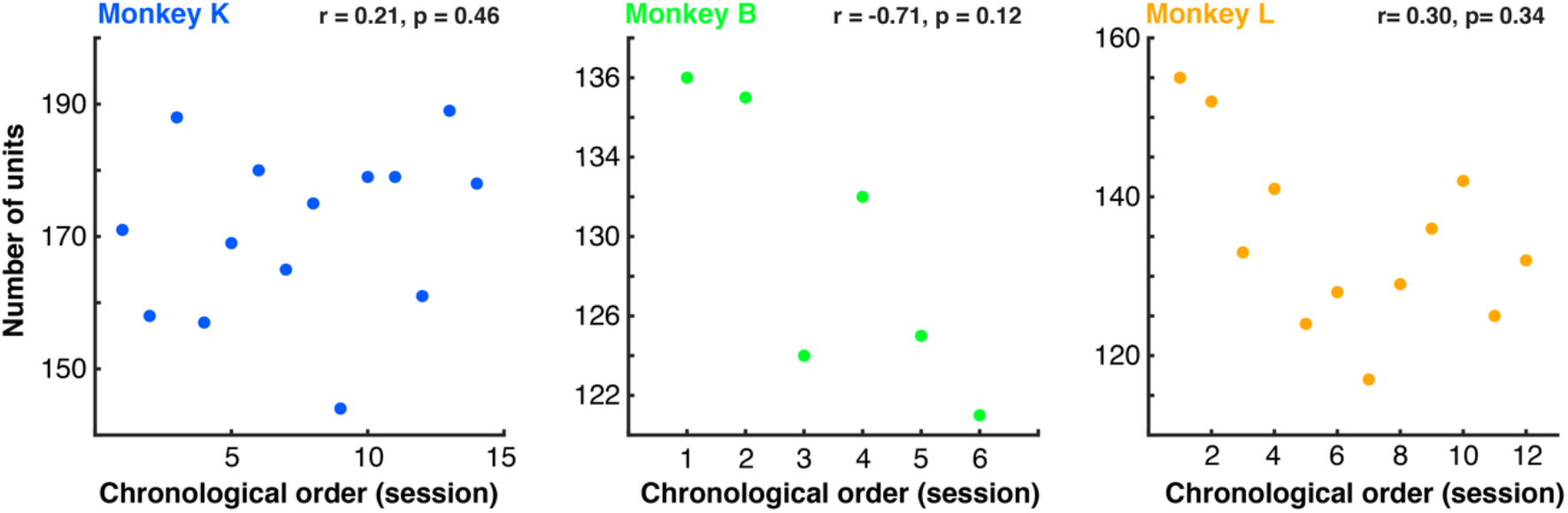
Number of recorded single and multiunits in every recording session, ordered chronologically. Number of recorded units was similar across the length of the experiment for all three monkeys.

**Supp. Figure 2.**
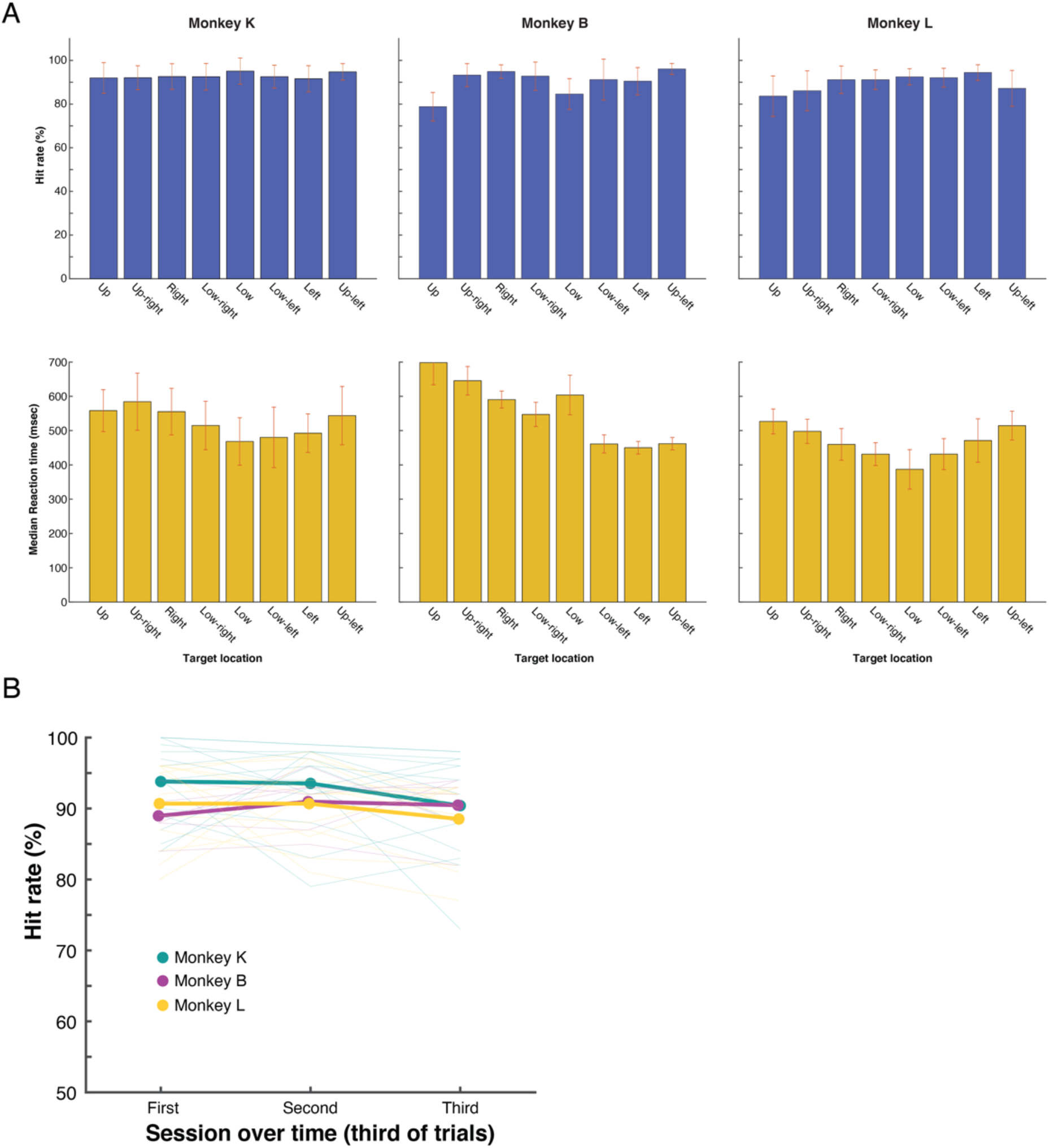
(A) Hit rate and reaction time as a function of target position (8 positions) during task performance for all three monkeys. (B) Hit rate over time during each recording session. Trials are split into thirds chronologically. Performance is stable across the length of all sessions.

**Supp. Figure 3.**
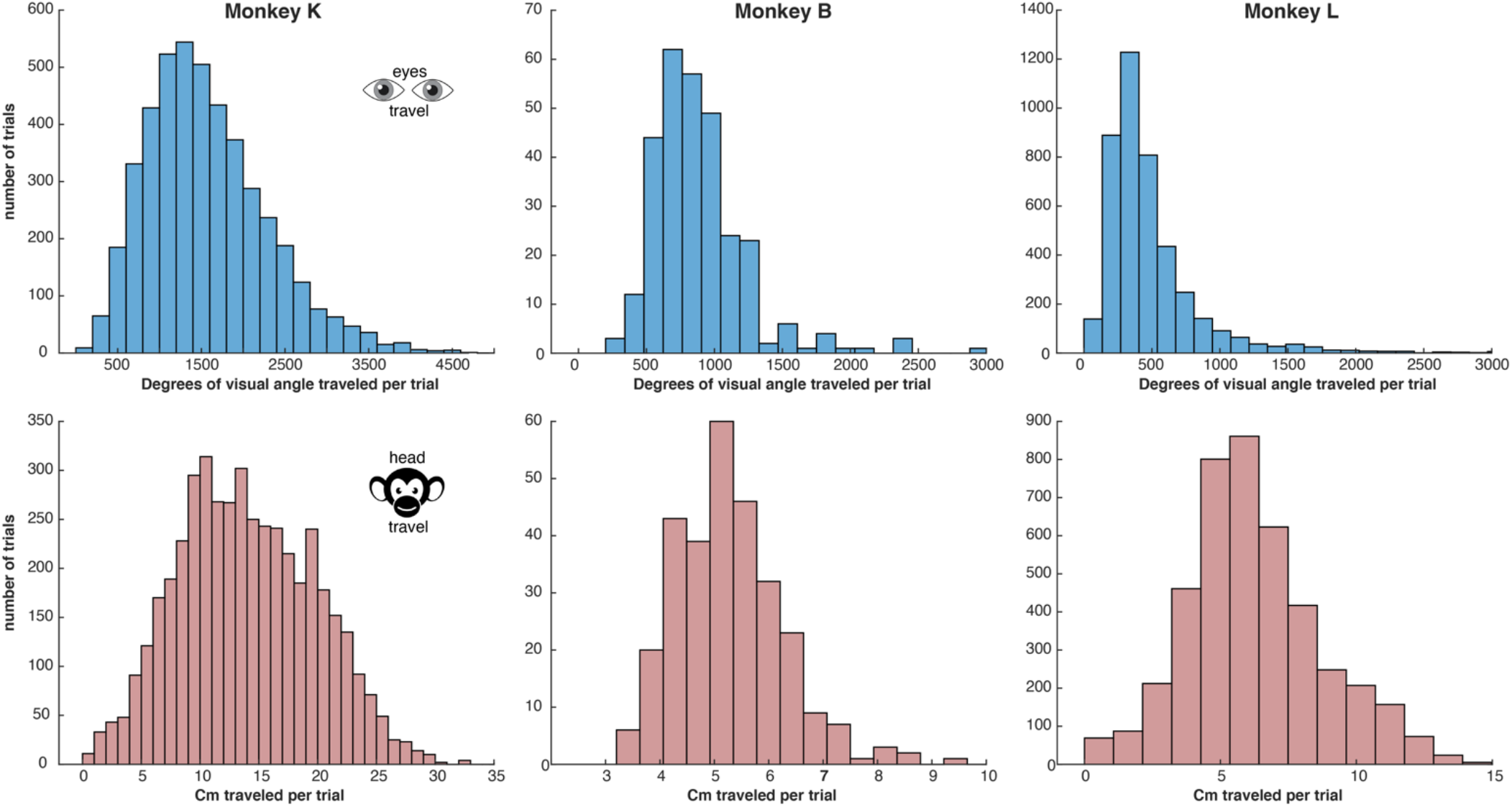
Quantification of movement variability from trial to trial for the eyes (top row) and the head (bottom row) for all three monkeys. Histograms represent the travel per trial; the total distance (in corresponding units) traveled by the head or the eyes over the course of one trial. There is a large variability in movement performed from trial to trial for these motor effectors.

**Supp. Figure 4.**
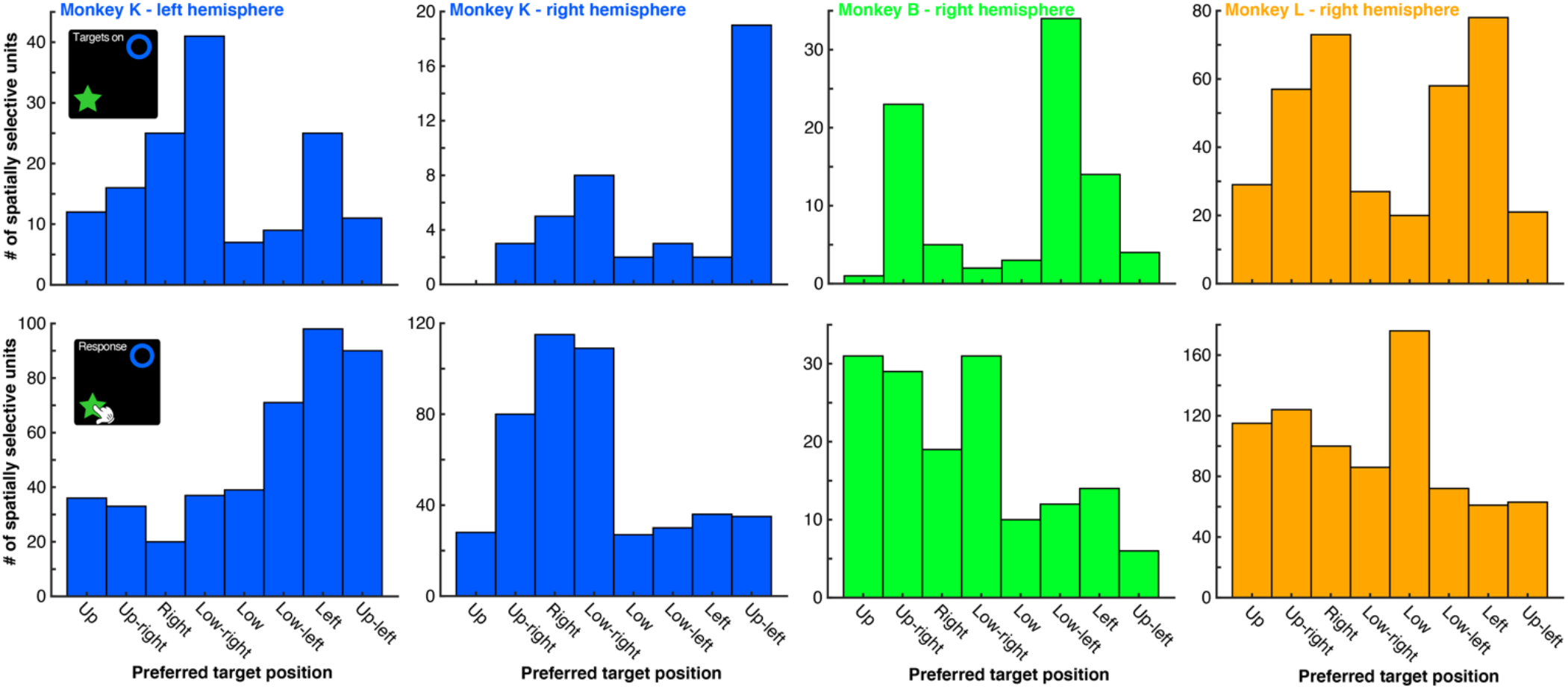
Spatial selectivities of all recorded neurons. Spatial selectivity of all recorded units during the target onset epoch (top) and the response selection epoch (bottom) for all three monkeys. The left and right hemisphere arrays are displayed separately for Monkey K. All arrays show a preponderance of neurons with ipsilateral selectivity during the response epoch, as opposed to the contralateral one seen with most eye movement paradigms.

**Supp. Figure 5.**
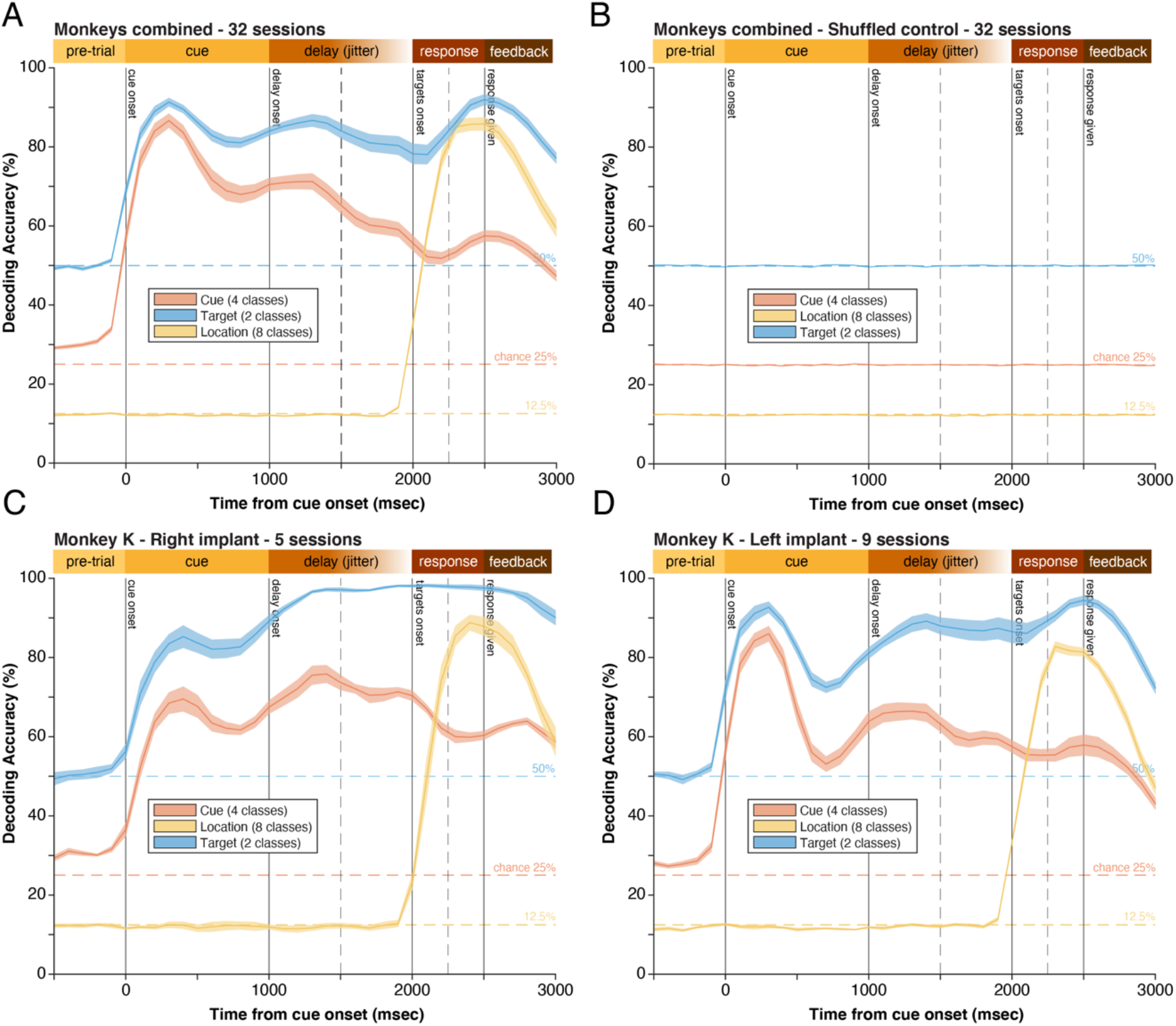
Decoding accuracies for combined monkeys. (A) Average decoding accuracies for instruction cue, target, and location across all three monkeys and sessions combined. Error bars are SEM. (B) Shuffled control whereby the labels used for training the decoder are randomly shuffled before training. Results fall to theoretical chance, expectedly. (C) Comparison of the decoding accuracies from the right hemisphere and the left hemisphere (D) in monkey K. Both Utah arrays were implanted in the same anatomical area (area 8A) in each hemisphere. Decoding results are comparable.

**Supp. Figure 6.**
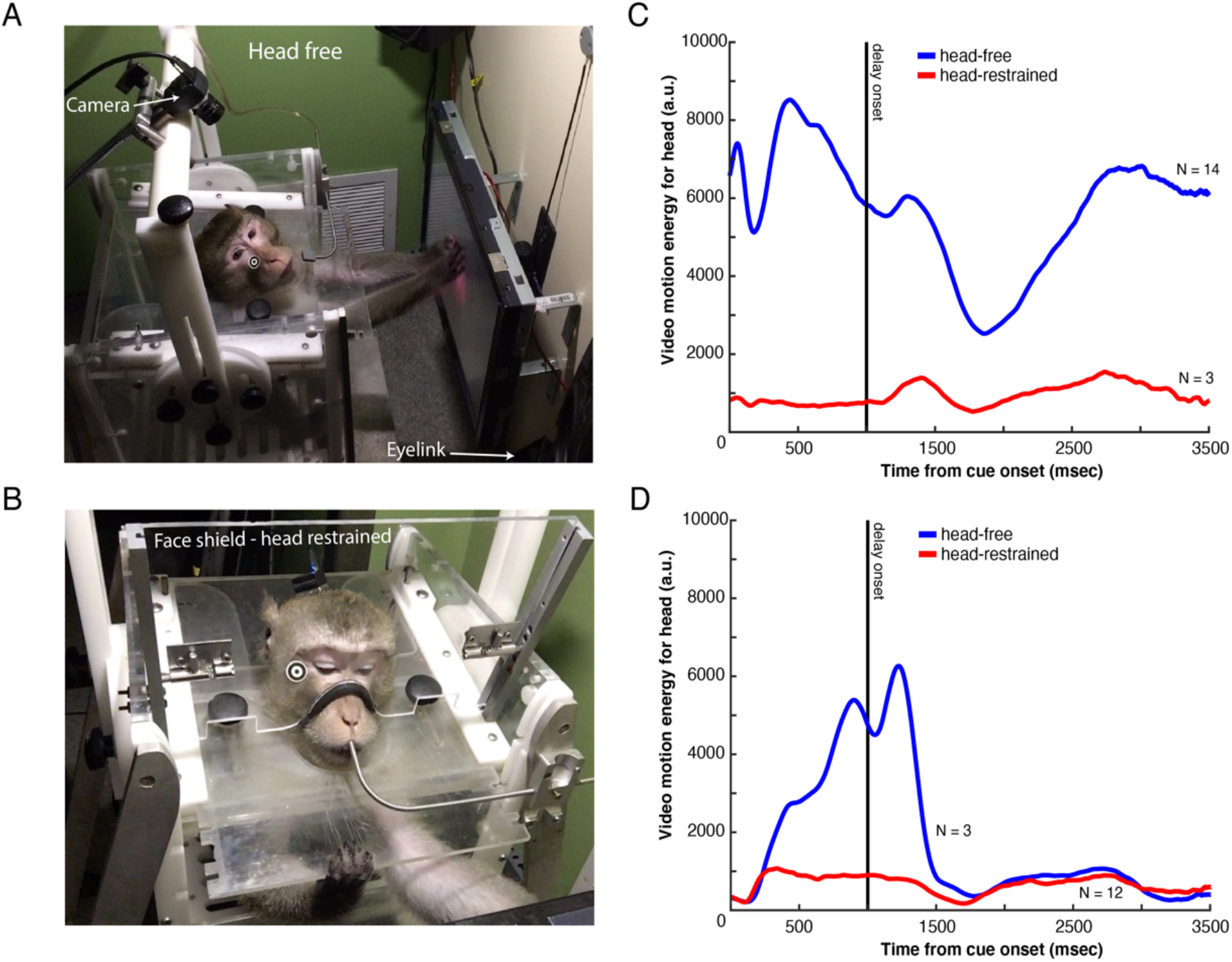
Head fixation and head motion energy. (A) Photograph of the experimental setup during an unrestricted, head-free recording session. (B) Photograph of the head-restraining device (aka. Face shield) used to stabilize the monkeys’ head during ‘head restrained’ recording sessions. (C) head motion energy (aka. movement) during head free recording sessions (blue) and head-restrained (red) recording sessions for monkey K. (D) Same as (C) but for monkey L.

**Supp. Figure 7.**
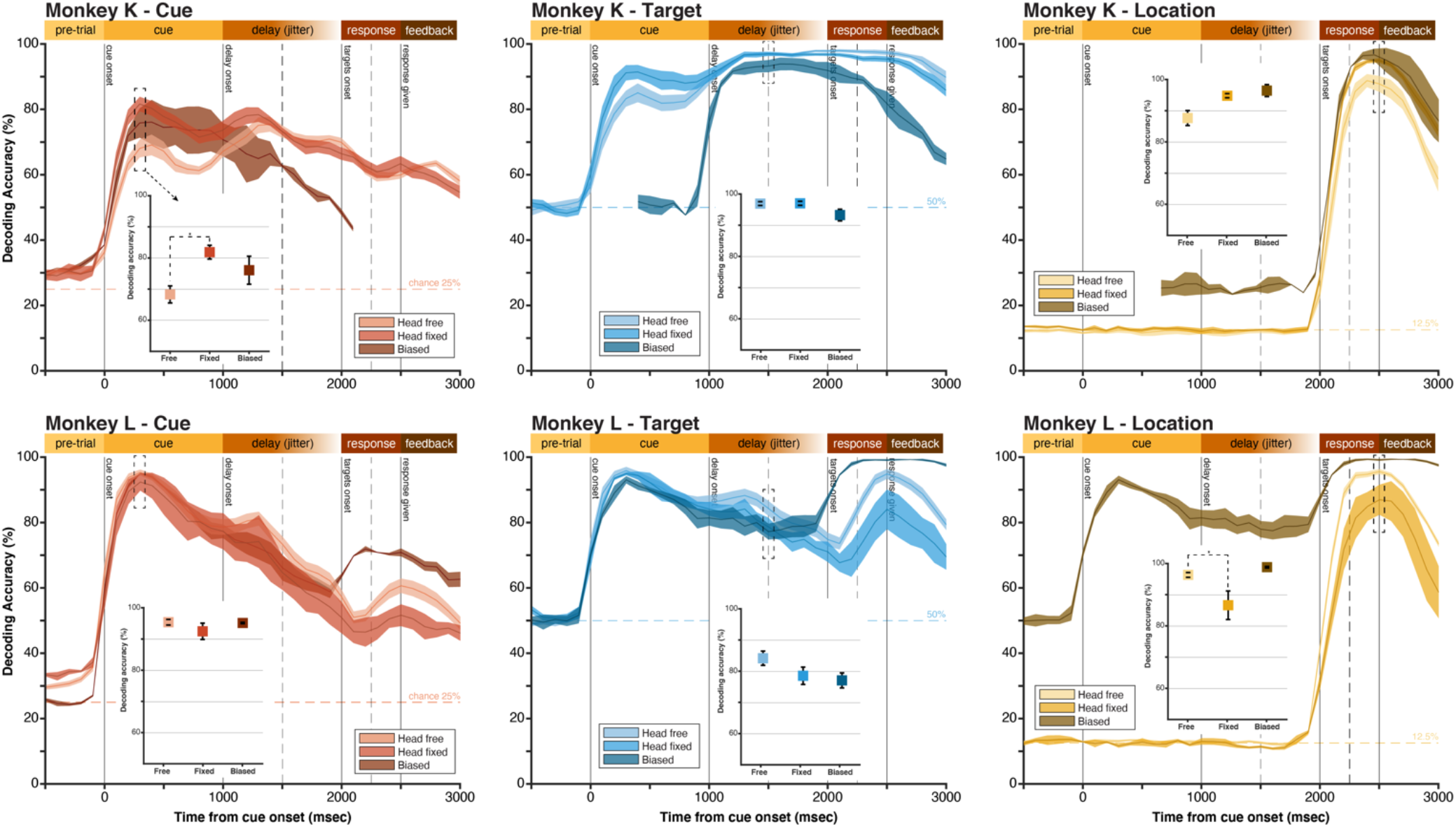
Decoding analyses in modified tasks. Decoding accuracies comparing variations of the behavioral task in which the monkey’s head is free to move (free), fixed to the primate chair (fixed), or spatially biased (biased). In monkey K, the biased task had a shorter trial length; decoding traces are aligned to key epochs instead. Insets represent decoding accuracies at key trial epochs. *: *P* < .05, FDR corrected. Error bars represent SEM.

**Supp. Figure 8.**
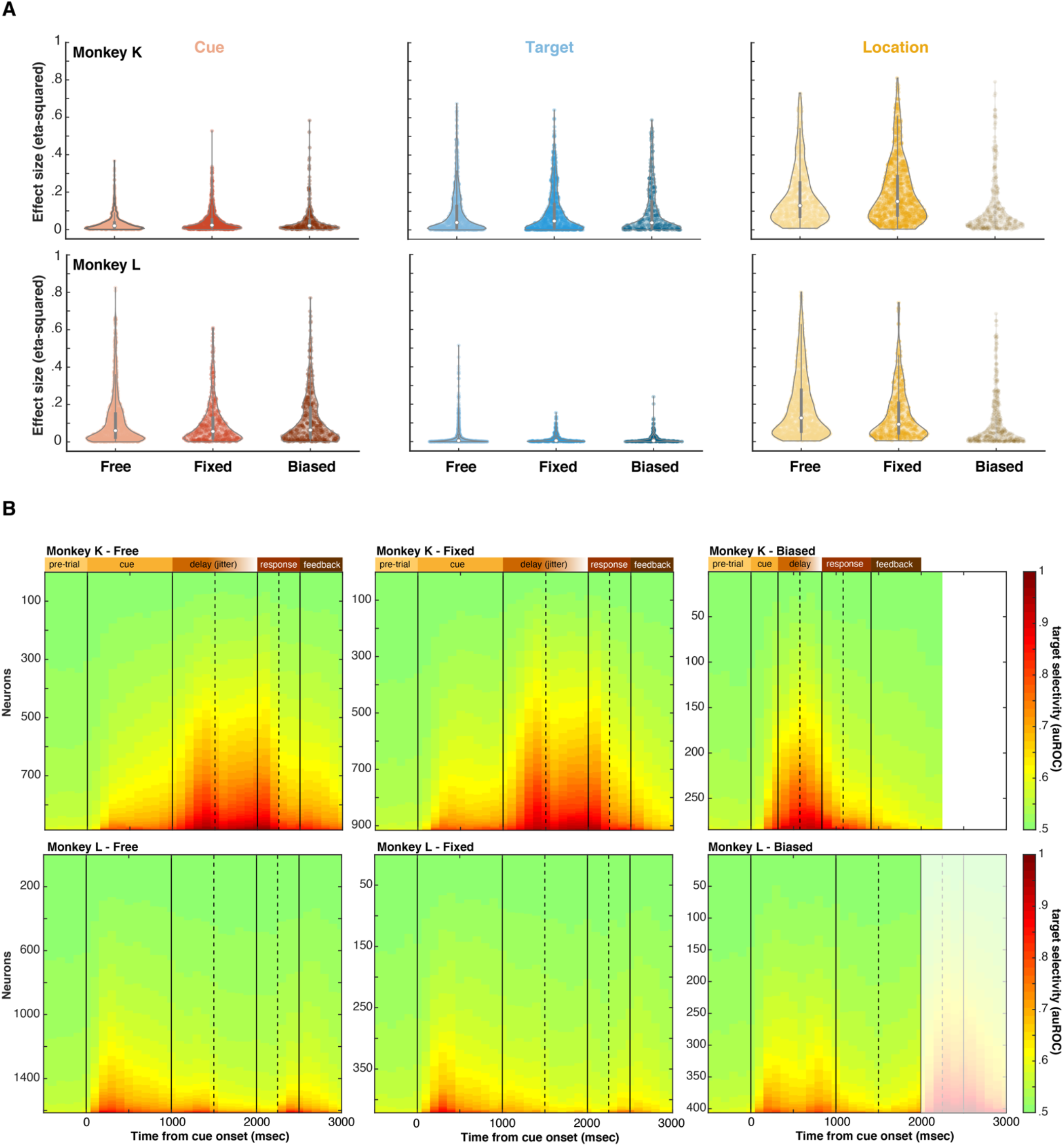
Analyses of tuning strength across behavioral manipulations. (A) Eta-squared metrics of effect size for the single neuron tuning to the cue (4 groups), target (2 groups), and location variables (8 groups) for all neurons recorded. Distributions are displayed as violin plots. Eta-squared metrics are biased by number of groups (higher with more groups). In the Biased tasks, there were fewer target locations (4 instead of 8) which explains lower values for the right-most plots (shaded). (B) Area under the receiving operating characteristic curve (auROC) for target selectivity during the entire trial for all neurons, separated by task. Neurons are ordered on the y-axis according to auROC value. For the “biased” task, trial length was shorter for monkey K, and location and target were confounded for monkey L during “response” (shaded area), explaining differences during these epochs specifically.

**Supp. Figure 9.**
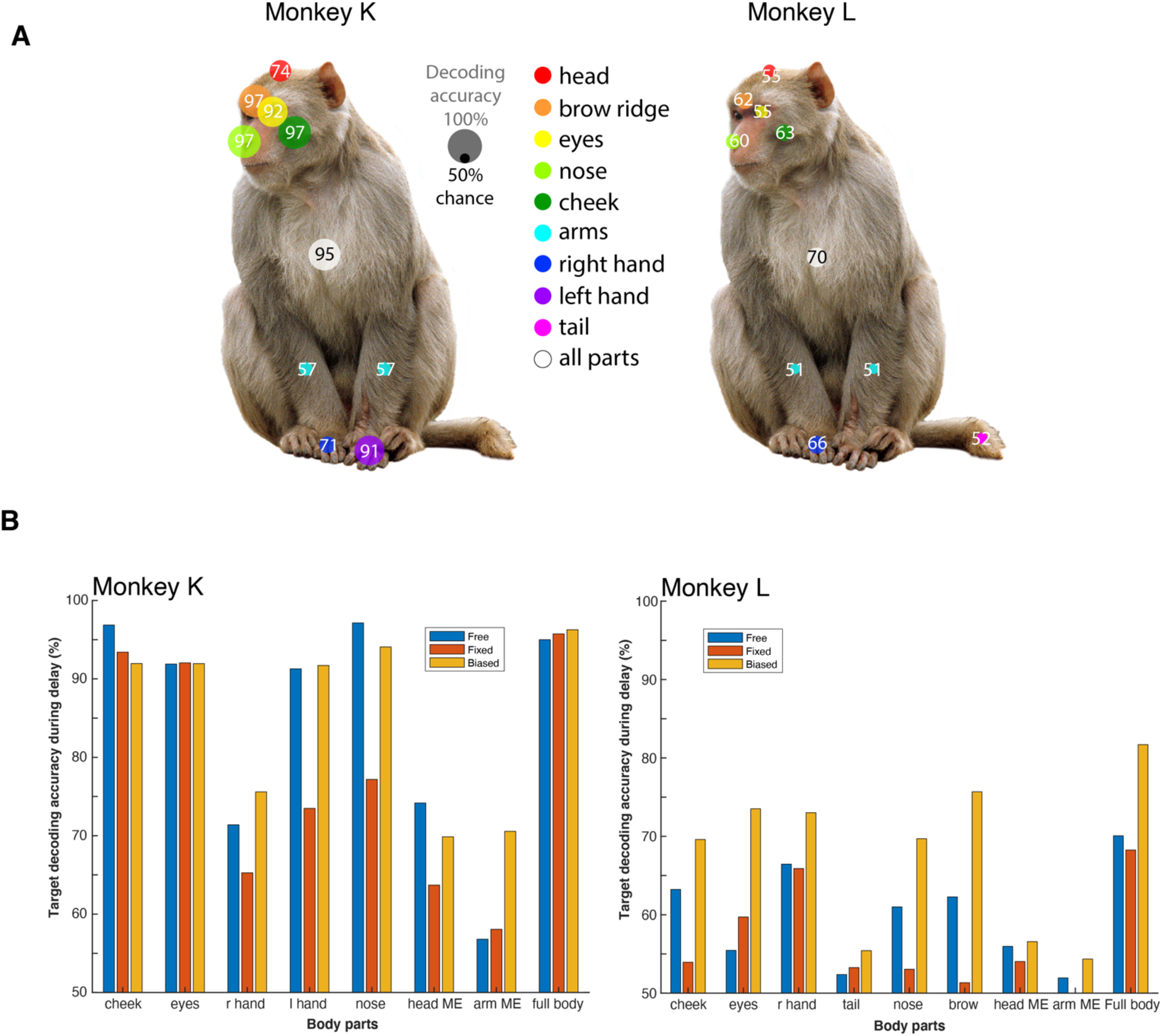
Decoding decisions from uninstructed movements. (A) Accuracy of a decoder predicting the decision (select star or circle) based only on uninstructed movement dynamics of motor effectors during the delay epoch, before the correct movement is known. Decoding accuracy of each effector is represented by circle size and overlaid number. Monkey K was left-handed and monkey L was right-handed. “All parts” represents a decoder where all motor effectors are included together as predictors. (B) Decoding accuracy of each motor effector during the delay when predicting upcoming decision, separated by behavioral conditions (Head-free, head-fixed, and biased tasks). “Full body” represents a decoder where all motor effectors are included together as predictors.

**Supp. Figure 10.**
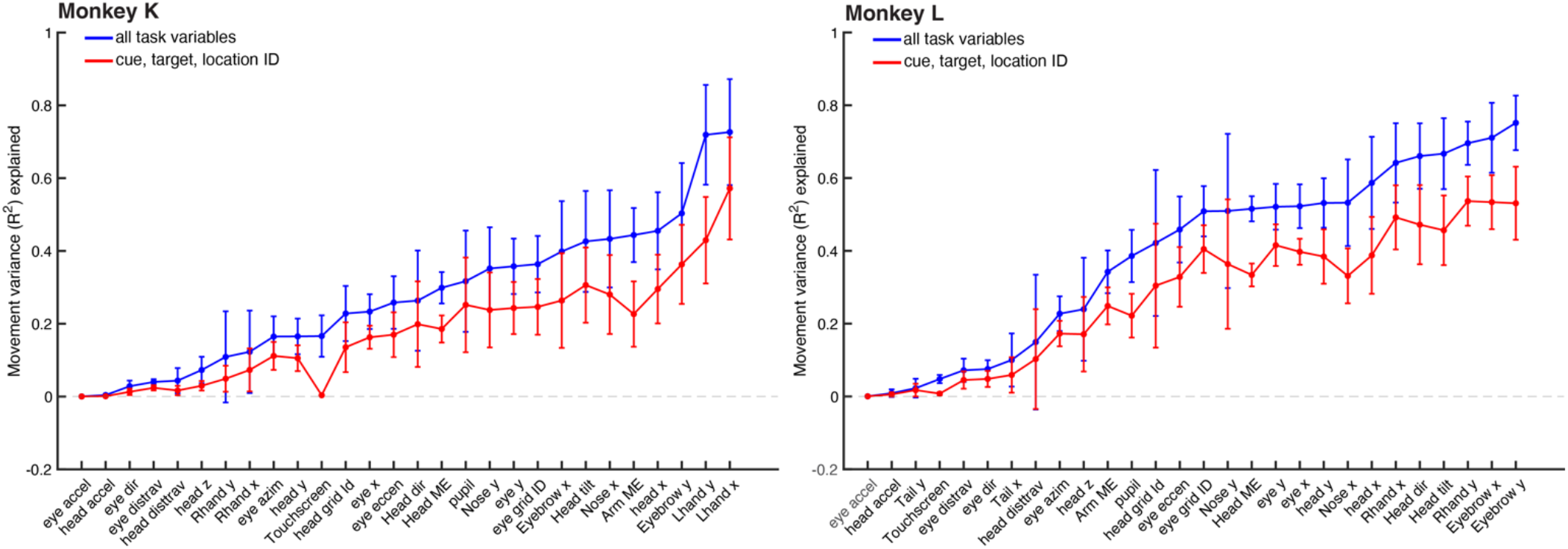
Movement variance explained by task variables. This analysis replicates the neural encoding analysis of Fig. 8, with the difference that single trial movement variance is predicted rather than neural population activity. Either all task variables (blue line) or only a subset of the main three task variables (cue, target, and location) are used to predict the position and movement of all motor effectors on a second-by-second basis (33 msec resolution). In both monkeys, task variables are highly predictive of body movements, indicating that those movements are highly correlated with key task variables.

**Supplementary Table 1.**
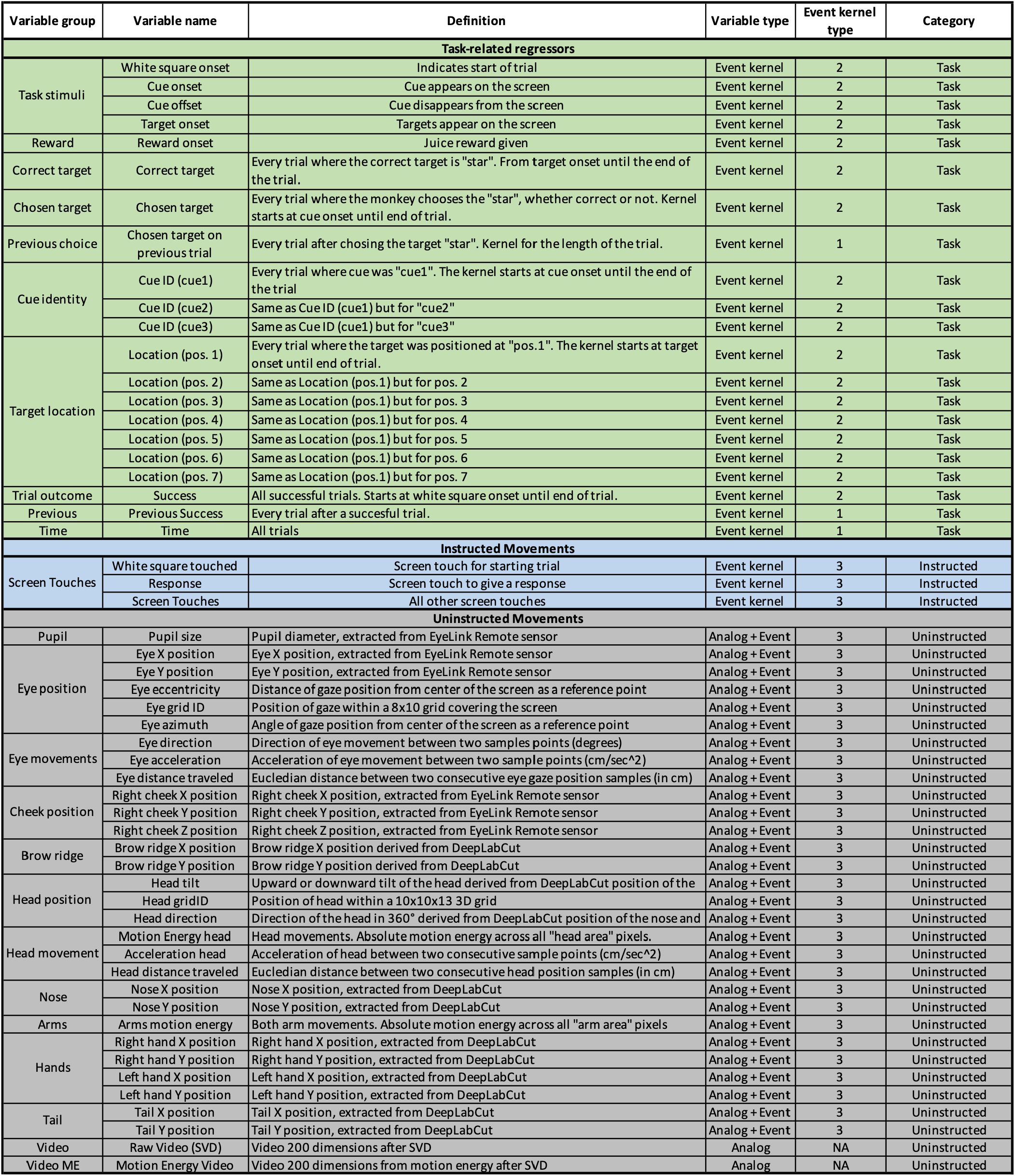
Model regressors. List of all variables inputted into the linear encoding model with their name, description, type of time varying kernel used (see Methods for details), and variable group category (task, instructed movement, uninstructed movement).

| Variable group | Variable name | Definition | Variable type | Event kernel type | Category |
| --- | --- | --- | --- | --- | --- |
| **Task-related regressors** |  |  |  |  |  |
| Task stimuli | White square onset | Indicates start of trial | Event kernel | 2 | Task |
|  | Cue onset | Cue appears on the screen | Event kernel | 2 | Task |
|  | Cue offset | Cue disappears from the screen | Event kernel | 2 | Task |
|  | Target onset | Targets appear on the screen | Event kernel | 2 | Task |
| Reward | Reward onset | Juice reward given | Event kernel | 2 | Task |
| Correct target | Correct target | Every trial where the correct target is "star". From target onset until the end of the trial. | Event kernel | 2 | Task |
| Chosen target | Chosen target | Every trial where the monkey chooses the "star", whether correct or not. Kernel starts at cue onset until end of trial. | Event kernel | 2 | Task |
| Previous choice | Chosen target on previous trial | Every trial after choosing the target "star". Kernel for the length of the trial. | Event kernel | 1 | Task |
| Cue identity | Cue ID (cue1) | Every trial where cue was "cue1". The kernel starts at cue onset until the end of the trial | Event kernel | 2 | Task |
|  | Cue ID (cue2) | Same as Cue ID (cue1) but for "cue2" | Event kernel | 2 | Task |
|  | Cue ID (cue3) | Same as Cue ID (cue1) but for "cue3" | Event kernel | 2 | Task |
| Target location | Location (pos. 1) | Every trial where the target was positioned at "pos.1". The kernel starts at target onset until end of trial. | Event kernel | 2 | Task |
|  | Location (pos. 2) | Same as Location (pos.1) but for pos. 2 | Event kernel | 2 | Task |
|  | Location (pos. 3) | Same as Location (pos.1) but for pos. 3 | Event kernel | 2 | Task |
|  | Location (pos. 4) | Same as Location (pos.1) but for pos. 4 | Event kernel | 2 | Task |
|  | Location (pos. 5) | Same as Location (pos.1) but for pos. 5 | Event kernel | 2 | Task |
|  | Location (pos. 6) | Same as Location (pos.1) but for pos. 6 | Event kernel | 2 | Task |
|  | Location (pos. 7) | Same as Location (pos.1) but for pos. 7 | Event kernel | 2 | Task |
| Trial outcome | Success | All successful trials. Starts at white square onset until end of trial. | Event kernel | 2 | Task |
| Previous | Previous Success | Every trial after a successful trial. | Event kernel | 1 | Task |
| Time | Time | All trials | Event kernel | 1 | Task |
| **Instructed Movements** |  |  |  |  |  |
| Screen Touches | White square touched | Screen touch for starting trial | Event kernel | 3 | Instructed |
|  | Response | Screen touch to give a response | Event kernel | 3 | Instructed |
|  | Screen Touches | All other screen touches | Event kernel | 3 | Instructed |
| **Uninstructed Movements** |  |  |  |  |  |
| Pupil | Pupil size | Pupil diameter, extracted from EyeLink Remote sensor | Analog + Event | 3 | Uninstructed |
| Eye position | Eye X position | Eye X position, extracted from EyeLink Remote sensor | Analog + Event | 3 | Uninstructed |
|  | Eye Y position | Eye Y position, extracted from EyeLink Remote sensor | Analog + Event | 3 | Uninstructed |
|  | Eye eccentricity | Distance of gaze position from center of the screen as a reference point | Analog + Event | 3 | Uninstructed |
|  | Eye grid ID | Position of gaze within a 8x10 grid covering the screen | Analog + Event | 3 | Uninstructed |
|  | Eye azimuth | Angle of gaze position from center of the screen as a reference point | Analog + Event | 3 | Uninstructed |
| Eye movements | Eye direction | Direction of eye movement between two samples points (degrees) | Analog + Event | 3 | Uninstructed |
|  | Eye acceleration | Acceleration of eye movement between two sample points (cm/sec<sup>2</sup>) | Analog + Event | 3 | Uninstructed |
|  | Eye distance traveled | Euclidian distance between two consecutive eye gaze position samples (in cm) | Analog + Event | 3 | Uninstructed |
| Cheek position | Right cheek X position | Right cheek X position, extracted from EyeLink Remote sensor | Analog + Event | 3 | Uninstructed |
|  | Right cheek Y position | Right cheek Y position, extracted from EyeLink Remote sensor | Analog + Event | 3 | Uninstructed |
|  | Right cheek Z position | Right cheek Z position, extracted from EyeLink Remote sensor | Analog + Event | 3 | Uninstructed |
| Brow ridge | Brow ridge X position | Brow ridge X position derived from DeepLabCut | Analog + Event | 3 | Uninstructed |
|  | Brow ridge Y position | Brow ridge Y position derived from DeepLabCut | Analog + Event | 3 | Uninstructed |
| Head position | Head tilt | Upward or downward tilt of the head derived from DeepLabCut position of the | Analog + Event | 3 | Uninstructed |
|  | Head gridID | Position of head within a 10x10x13 3D grid | Analog + Event | 3 | Uninstructed |
|  | Head direction | Direction of the head in 360° derived from DeepLabCut position of the nose and | Analog + Event | 3 | Uninstructed |
| Head movement | Motion Energy head | Head movements. Absolute motion energy across all "head area" pixels. | Analog + Event | 3 | Uninstructed |
|  | Acceleration head | Acceleration of head between two consecutive sample points (cm/sec<sup>2</sup>) | Analog + Event | 3 | Uninstructed |
|  | Head distance traveled | Euclidian distance between two consecutive head position samples (in cm) | Analog + Event | 3 | Uninstructed |
| Nose | Nose X position | Nose X position, extracted from DeepLabCut | Analog + Event | 3 | Uninstructed |
|  | Nose Y position | Nose Y position, extracted from DeepLabCut | Analog + Event | 3 | Uninstructed |
| Arms | Arms motion energy | Both arm movements. Absolute motion energy across all "arm area" pixels | Analog + Event | 3 | Uninstructed |
| Hands | Right hand X position | Right hand X position, extracted from DeepLabCut | Analog + Event | 3 | Uninstructed |
|  | Right hand Y position | Right hand Y position, extracted from DeepLabCut | Analog + Event | 3 | Uninstructed |
|  | Left hand X position | Left hand X position, extracted from DeepLabCut | Analog + Event | 3 | Uninstructed |
|  | Left hand Y position | Left hand Y position, extracted from DeepLabCut | Analog + Event | 3 | Uninstructed |
| Tail | Tail X position | Tail X position, extracted from DeepLabCut | Analog + Event | 3 | Uninstructed |
|  | Tail Y position | Tail Y position, extracted from DeepLabCut | Analog + Event | 3 | Uninstructed |
| Video | Raw Video (SVD) | Video 200 dimensions after SVD | Analog | NA | Uninstructed |
| Video ME | Motion Energy Video | Video 200 dimensions from motion energy after SVD | Analog | NA | Uninstructed |

## Acknowledgements

We thank S. Musall, M.T. Kaufman, and A.K. Churchland in providing assistance to replicate the methodology used in Musall et al. *Nat. Neurosci.* 2019. Financial support was received from the Canadian Institutes of Health Research Fellowship Award (S.T.), the NIH (R.D. 5R01DC017690-02), and a Natural Sciences and Engineering Research Council of Canada grant to M.P.

## Author contributions

S.T., C.T., and M.P. designed the experiments. S.T., C.T., and J.I. trained the animals and recorded the data. S.T., C.T., and R.D. analyzed the data. S.T., C.T., and M.P. wrote the article with assistance from J.I. and R.D.

